# Identification of the Metaphyseal Skeletal Stem Cell

**DOI:** 10.1101/2022.09.08.506930

**Authors:** Guan Yang, Qi He, Xiaoxiao Guo, Rong-Yu Li, Jingting Lin, Wanyu Tao, Wenjia Liu, Huisang Lin, Mingchuan Tang, Shilai Xing, Yini Qi, Yanli Peng, Lei Dong, Jingdong Han, Bin Zhou, Yan Teng, Xiao Yang

## Abstract

Identification of novel regional skeletal stem cells (SSCs) will provide a new cellular paradigm for bone physiology and dysfunction. Several populations of SSCs have been identified at distinct skeletal sites. However, a bona fide SSC population has not yet been formally characterized in the bone marrow. Here, we identify a metaphyseal SSCs (mpSSCs) population whose transcriptional landscape is distinct from other bone mesenchymal stromal cells (bMSCs) in the bone marrow. These mpSSCs emerge at the postnatal stage and reside just underneath the growth plate, consistent with the fact that these mpSSCs are exclusively derived from hypertrophic chondrocytes (HCs). These mpSSCs possess SSC properties such as self-renewal and multipotency *in vitro* and *in vivo*, stand at the top of the HC de-differentiation path, and produce most HC progeny. Genetic block of the conversion from HCs to mpSSCs significantly compromises trabecular bone formation and bone regeneration. Thus, metaphysis houses a unique HC-derived SSC population, which is a major source of osteoblasts and bMSCs supporting postnatal trabecular bone formation.

## INTRODUCTION

Skeletal stem cells (SSCs) stand at the apex of the skeletal lineage, with the properties of self-renewal and generating osteoblast, chondrocytes, adipocytes, and bone mesenchymal stromal cells (bMSCs), thus participating in growth, maintenance, and repair of skeleton (Ambrosi et al., 2019; Serowoky et al., 2020). Cell surface markers such as Pdgfrb, Pdgfra, and Sca-1 have long been used for isolating murine bMSC (Koide et al., 2007; Morikawa et al., 2009), but the surface markers configuration for SSCs have been identified in recent years. A pioneering study using a panel of cell surface markers CD200^+^CD51^+^CD90^−^Ly51^−^CD105^−^ defines murine SSCs in the developing bone, which can produce eight defined and restricted stromal progenitor cell types when transplanted into the kidney capsules of immunodeficient mice (Chan et al., 2015). Although it is not tractable to examine the exact anatomic location of these functional SSCs defined by the multiple surface markers, subsequent genetic lineage tracing studies have demonstrated that they relate to several SSC populations within the periosteum, resting chondrocytes, articular surfaces, and the mandible (Debnath et al., 2018; Mizuhashi et al., 2018; Murphy et al., 2020; Ransom et al., 2018). However, whether the CD200^+^CD51^+^ functional SSCs exist in the bone marrow is not clear.

Bone marrow is a composite of diverse cell types, including cells of osseous, adipogenic, stromal, endothelial, and hematopoietic lineages (Baryawno et al., 2019; Kfoury and Scadden, 2015; Tikhonova et al., 2019). Among these cells, bMSC is a predominant cell type and has important roles in generating skeletal lineage cells and supporting hematopoiesis. BMSCs are actually a heterogenous mixture of different cell subpopulations with distinct markers, spatiotemporal characteristics, and functions. BMSCs have varying progenitor properties, but this may be attributed to a rare fraction of bMSCs -- bona fide SSCs that emerge during development and have a multilineage differentiation capacity. A recent report showed that metaphyseal bMSCs have greater potential for growth and multipotential differentiation than diaphyseal bMSCs, raising the possibility that if SSCs exist under the growth plate, they may reside in the metaphysis (Sivaraj et al., 2021). A previous report argues that murine metaphysis harbors a SSC population labeled by Bmp antagonist gremlin 1 (Grem1) (Worthley et al., 2015). The important evidence to support Grem1^+^ cells as endogenous SSCs is lineage tracing showing that Grem1^+^ cells can give rise to growth plate chondrocytes, osteoblasts, and bMSCs. However, because Grem1 can concurrently label a considerable number of chondrocytes, the multilineage differentiation potential of the alleged Grem1^+^ SSCs may reflect the well-established notion that hypertrophic chondrocytes (HCs) transdifferentiate into osteoblasts and bMSCs, challenging Grem1^+^ cells to be true SSCs. Besides Grem1^+^ cells, metaphysis also houses Gli1^+^ and Osterix^+^ progenitors, which can produce trabecular osteoblasts and bMSCs (Mizoguchi et al., 2014; Shi et al., 2017). However, a bona fide metaphyseal SSC population has not yet been formally identified, let alone its relation to these previously identified populations.

Here, through mapping the *in situ* location of CD200^+^CD51^+^CD90^−^Ly51^−^CD105^−^ SSCs and their enrichment in the single-cell transcriptome of the bulk bone marrow, we identify a putative metaphyseal SSCs with a distinct transcriptional landscape. These mpSSCs emerge at the postnatal stage and are restricted to immediately underneath the growth plate, consistent with the fact that mpSSCs are the first cells that HCs differentiate into when entering the metaphysis. The mpSSCs meet the SSC criteria, such as the self-renewal and multilineage differentiation and serve as a major source of trabecular osteoblasts as well as bMSCs in the lower metaphyseal and diaphyseal bone marrow. Hepatocyte growth factor-regulated tyrosine kinase substrate (Hgs) is highly expressed in a subpopulation of HCs and drives the conversion of HCs to mpSSCs. A reduced mpSSC pool resulting from loss of Hgs compromises trabecular bone formation and bone regeneration.

## RESULTS

### A Population of CD200^+^CD51^+^ Murine SSCs Are Confined to Metaphysis

We first determined the *in situ* localization of CD200^+^CD51^+^Thy1^−^Ly51^−^CD105^−^CD45^−^ Ter119^−^CD31^−^ murine SSCs (Chan et al., 2015) in the femurs on postnatal day 8 (P8) using multicolor immunofluorescence (IF) staining (Figures 1A and 1B). Consistent with the previous reports, these SSCs were enriched in periosteum and articular surface (Figure 1B). Interestingly, these SSCs were also observed at other anatomic sites with robust bone formation, including the secondary ossification center, cortical bone endosteum, metaphyseal trabeculae (Figure 1B). At metaphysis, CD200^+^CD51^+^Thy1^−^Ly51^−^CD105^−^CD45^−^ Ter119^−^ CD31^−^ cells resided right underneath the growth plate (Figure 1B).

**Figure 1.**
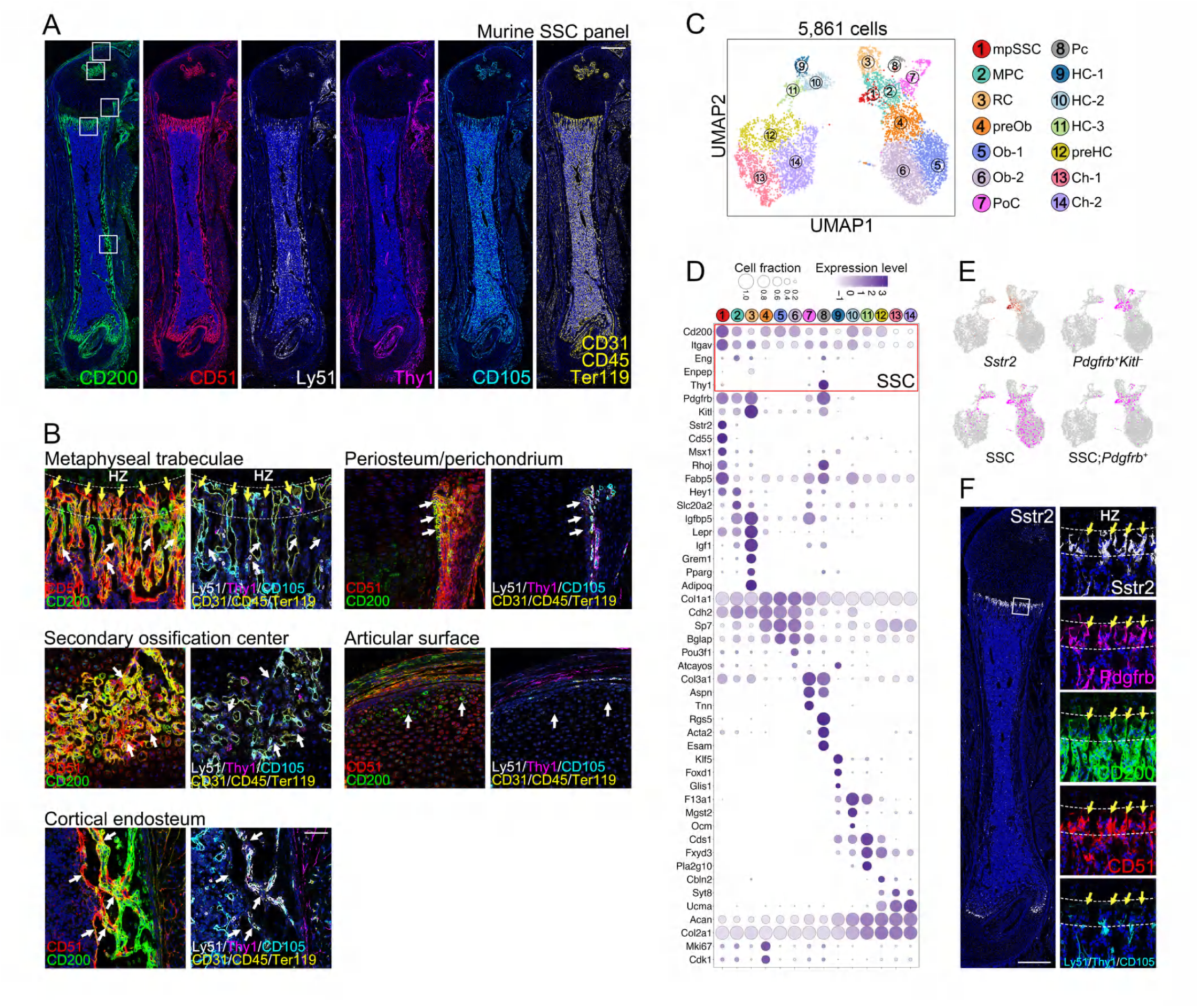
Identification of a Metaphyseal Murine SSC Population. (A, B) Representative multicolor immunofluorescent staining shows the expression of murine SSC panel in P8 femur section. White framed areas are shown in (B) at higher magnifications, which demonstrate CD200^+^CD51^+^Thy1^−^Ly51^−^CD105^−^CD45^−^Ter119^−^ CD31^−^ mSSCs at the metaphyseal trabeculae, the periosteum and perichondrium, the secondary ossification center, the articular surface, and the cortical endosteum (white arrows). The yellow arrows pinpoint a cluster of SSCs at the upper metaphyseal region right beneath the hypertrophic zone (HZ) (framed by dashed lines). Scale bar, 500 μm. (C) ScRNA-seq of 5,861 skeletal cells from P7.5 hindlimb bones is visualized by uniform manifold approximation and projection (UMAP) (also see Methods and Figure S1). They are unbiasedly clustered into 14 clusters that include bMSCs (cluster 1–3), OLCs (cluster 4–6), periosteal cells (cluster 7), pericytes (cluster 8), and chondrocytes (cluster 9–14). (D) Dot plot shows the expression level of selected cluster-enriched genes in 14 clusters. Dot size indicates the cell fraction of each cluster that expresses listed genes. Purplish grey color intensity indicates scaled average expression level. Black-coded frames indicate murine SSC panel genes *Cd200* (CD200), *Itgav* (CD51), *Eng* (CD105), *Enpep* (Ly51), and Thy1. (E) Enriched gene signatures of cluster 1 (mpSSC). Genes with count >1 are identified as positive expressed genes (+); count ≤0 are identified as negative (–). Cells filtered by combined gene expression patterns are labeled with magenta dots and are superimposed on the UMAP plot. (F) Representative multicolor immunofluorescent staining shows the expression of mpSSC markers in P7.5 femur section. White framed areas are shown adjacently at higher magnifications. The upper metaphyseal region right beneath the hypertrophic zone (HZ) is framed by dashed lines. Yellow arrows indicated mpSSCs that are labeled by Sstr2^+^ or SSC;Pdgfrb^+^. Scale bar, 500 μm.

We hypothesized that these putative SSCs at metaphysis would represent a distinct population in the bone marrow. ScRNA-seq analysis using the 10× Genomics Chromium platform was performed to characterize all skeletal cells in the P7.5 tibia and femur (Figure S1). ScRNA-seq transcriptomes of 5,861 skeletal cells were visualized using a uniform manifold approximation and projection (UMAP). Mapping of historically defined marker genes onto the UMAP representation revealed 14 clusters, generally falling into three major subpopulations according to their gene signature: 1) stromal cells expressing *Pdgfrb*, including cluster 1 (metaphyseal skeletal stem cell, mpSSC), cluster 2 (mesenchymal progenitor cell, MPC), cluster 3 (reticular cell, RC), cluster 7 (periosteal cell, PoC), and cluster 8 (pericyte, Pc); 2) Osteo-lineage cells (OLCs) marked by *Col1a1*, including clusters 4 (pre-osteoblast, preOb), cluster 5 and 6 (osteoblast, Ob-1 and -2); 3) chondrocytes and HCs marked by *Acan*, including clusters 9–11 (HC-1, -2, and -3), cluster 12 (preHC), and cluster 13 and 14 (chondrocytes, Ch-1 and -2) (Figures 1C and 1D).

Next, scRNA-seq data were filtered by the SSC gene signature *CD200*^+^*Itgav*^+^*Thy1*^−^*Enpep*^−^*Eng*^−^ (CD200^+^CD51^+^Thy1^−^Ly51^−^CD105^−^). Among all 14 clusters, cluster 1 seemed to share the most enriched gene expression patterns with SSCs (Figures 1D and 1E). This population was separated from other skeletal cell clusters by somatostatin receptor 2 (*Sstr2*), as well as *Pdgfrb^high^Kitl^low^*, *Cd55*, *Msx1*, and *Rhoj* (Figures 1D and 1E). Notably, a single marker Sstr2 could specifically label this subpopulation throughout all clusters (Figure 1D and 1E). Multicolor IF staining revealed that Sstr2^+^ cells resided in the upper trabecular zone and right beneath the hypertrophic cartilage, expressed the general stromal cells marker Pdgfrb, and extensively overlapped with the CD200^+^CD51^+^Thy1^−^Ly51^−^CD105^−^ murine SSCs (Figures 1E and 1D, yellow arrows).

Taken together, these above data suggest that a putative Pdgfrb^+^ population of murine SSCs, signatured by a single marker Sstr2, resides at metaphysis and has a distinct transcriptional profile within the bMSC populations.

### MpSSCs Are Derived from HCs

Previous genetic lineage tracing studies have demonstrated that most HCs can survive at the chondro-osseous junction and further convert into OLCs, adipocytes, or bSMCs (Giovannone et al., 2019; Long et al., 2022; Mizuhashi et al., 2019; Mizuhashi et al., 2018; Park et al., 2015; Shu et al., 2021; Yang et al., 2014a; Yang et al., 2014b; Zhou et al., 2014b). Given that most HC progeny are enriched in the metaphysis, it is conceivable that HC is the cellular source of the putative SSC at the metaphysis. scRNA-seq analysis of ZsGreen^+^ cells from tibias and femurs of P7.5 *Col10a1- Cre;ROSA26^LSL-ZsGreen^*mice (*Col10a1^ZsGreen^*) was performed. 1,998 ZsGreen^+^ cells were visualized using UMAP and 11 clusters were divided into three major subpopulations: 1) HCs marked by *Col10a1* (clusters 1, 2, 3); 2) bMSCs expressing *Pdgfrb* (clusters 4, 5, 6, and part of cluster 7); 3) OLCs marked by *Col1a1* (clusters 8, 9, 10, 11, and part of cluster 7) (Figures 2A and 2B). Notably, cluster 4 (mpSSC) was marked by Sstr2 (Figures 2B and 2C, white arrows) as well as *Cd55*, *Pdgfrb^high^Kitl^low^*, *Msx1*, and *Rhoj* (Figures S2A and S2B). The transcriptional signature of Sstr2^+^ cluster 4 was similar to that of Sstr2^+^ cluster 1 of the bulk scRNA-seq (Figures 1D and 1E), indicating that they might be the overlapping cluster. Indeed, multiple IF staining of *Col10a1-Cre;ROSA26^LSL-tdTomato^;Kitl^GFP^* mice (*Col10a1^tdTomato^;Kitl^GFP^*) revealed that all Sstr2^+^ cells are positive for tdTomato, suggesting HCs as the single source for the Sstr2^+^ putative SSCs (Figure 2C). To further corroborate it, we further generated an *Ihh-mKate2^tomm20^-Dre* knock-in mouse line in which Dre recombinase is restrictively expressed in pre-HCs (Li et al., 2022) (Figures S3A and S3B). Again, IF analysis of femurs of P8 *Ihh-mKate2^tomm20^-Dre;ROSA26^RSR-tdTomato^*mice confirmed that HCs contribute to multiple skeletal lineage under the growth plate, and that Sstr2^+^ cells are exclusively derived from HCs (Figure S3C).

**Figure 2.**
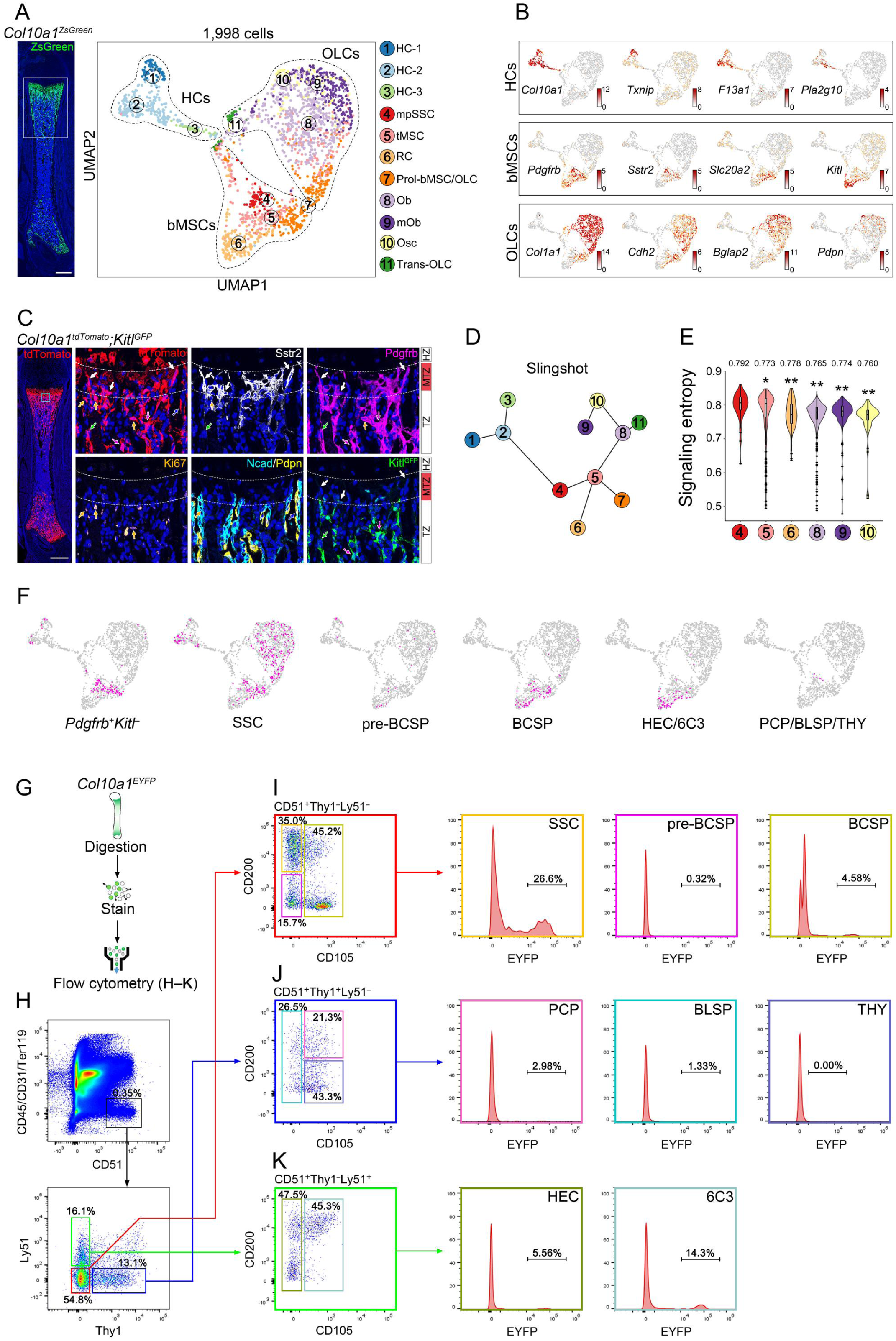
MpSSCs Are Derived from HCs. (A) The framed parts of hindlimb bones of P7.5 *Col10a1^ZsGreen^* mice were used for scRNA-seq of hypertrophic chondrocytes (HCs) and their descendants, and UMAP shows 11 clusters divided into three major subpopulations including HCs, bMSCs, and OLCs. Scale bar, 500 μm. (B) Selected cluster-enriched genes are superimposed on the UMAP plot shown in (A). Crimson color intensity indicates the Log2 expression level of genes. (C) Multicolor IF staining locates the main clusters of HC-derived bMSCs and OLCs by the expression of cluster-enriched genes shown in (B) in the femur of P7 *Col10a1^tdTomato^;Kitl^GFP^*mice. The cyan-framed areas are shown adjacently at higher magnifications. Dashed lines frame the upper metaphyseal region right beneath the hypertrophic zone. White arrows, tdTomato^+^Sstr2^+^Pdgfrb^high^Kitl^low/–^ mpSSCs; green arrows, tdTomato^+^Sstr2^−^Pdgfrb^+^Kitl^low^ tMSCs; magenta arrows, tdTomato^+^Pdgfrb^+^Kitl^high^ RCs; blue arrow, tdTomato^+^Ncad^+^/Pdpn^+^ OLCs; gold arrows, Tomato^+^Ki67^+^ proliferating mpSSCs, tMSCs and OLCs. HZ, hypertrophic zone; MTZ, metaphyseal trabecular zone; TZ, trabecular zone. Scale bar, 500 μm. (D) Differentiation trajectory was inferred by Slingshot which starts with the HC-1 cluster. (E) Signaling entropy analysis indicates mpSSCs (cluster 4) has a higher differentiation potency than other bMSCs and OLCs. Significant differences between mpSSC and other HC progeny clusters were analyzed using the t-test. *p-value < 0.05; **p-value < 0.01. (F) Identification of HC-derived mpSSCs, SSCs, and other lineage progenitor cells based on the gene expression profiles. Genes with count >1 are identified as positive expressed genes (+); count ≤0 are identified as negative (–). Cells filtered by combined gene expression patterns are labeled with magenta dots and are superimposed on the UMAP plot. (G) Scheme showing the phenotypic characterization of HC-derived murine SSC lineages. (H–K) Representative flow cytometry plots of cells digested from hindlimb bones of P7–P10 *Col10a1^EYFP^* mice, showing the putative contribution of HC-derived progeny to the compartments of the whole murine SSC lineages.

The specific location of Sstr2^+^ cells raised a possibility that the Sstr2^+^ putative SSCs may be originated from direct conversion of HCs. Pseudotime analyses including Slingshot and SCENT algorithm were used to infer the hierarchical relationships between all clusters of HCs progeny (Street et al., 2018; Teschendorff and Enver, 2017). Slingshot revealed a clear transcriptional continuum from HCs (clusters 1-3) to mpSSCs (cluster 4) and then tMSCs (trabecular mesenchymal stromal cells, cluster 5), followed by branching into RCs, Prol-bMSCs/OLCs, and Obs (clusters 6-8) (Figures 2D). SCENT algorithm, which characterizes the developmental plasticity of multipotent stem and progenitor cells by calculating the signaling entropy rate based on gene and protein signaling diversity, revealed that cluster 4 displays the highest entropy among all HC progeny clusters, indicating that mpSSCs have higher differentiation potential than other bMSCs and OLCs (Figure 2E). Therefore, HCs seem to directly de-differentiate into *Sstr2^+^* or *Pdgfrb^high^Kitl^low^*cells, which may be a new SSC population standing at the top of the hierarchy along the HC dedifferentiation trajectory.

Finally, we interrogated the correlation between HC-derived *Sstr2^+^*or *Pdgfrb^high^Kitl^low^* cells and CD200^+^CD51^+^Thy1^−^Ly51^−^CD105^−^ SSCs. As expected, cluster 4 (mpSSC) highly expressed the signature genes of SSCs (CD200^+^CD51^+^Thy1^−^Ly51^−^CD105^−^), while the clusters 5 and 6 (tMSC/Prol-bMSC/OLC and RC) subpopulations, the putative downstream progeny of cluster 4, expressed the transcripts associated with progenitor BCSPs (short for bone, cartilage, and stromal progenitors, sorted by CD51^+^Thy1^−^Ly51^−^CD105^+^) and HECs/6C3s (belonging to stromal hepatic leukemia factor-expressing cells, sorted by CD51^+^Thy1^−^Ly51^+^CD105^+/–^), respectively (Figure 2F). Because both BCSPs and HECs/6C3s are proposed to be the descendants of CD200^+^CD51^+^Thy1^−^Ly51^−^CD105^−^ SSCs, the transcriptome of HC progeny suggested that the HC dedifferentiation path would follow the hierarchy of CD200^+^CD51^+^ SSCs as previously described *in vitro*. Furthermore, the whole bone flow cytometry analysis revealed that HC descendants constituted 25.8% ± 5.6% of murine SSCs, 5.9% ± 1.8% BCSPs, 6.1% ± 2.0% CD51^+^Thy1^−^Ly51^+^ CD105^−^ HECs, and 14.4% ± 1.9% CD51^+^Thy1^−^Ly51^+^CD105^+^ 6C3s, while contributed relatively less to pre-BCSPs (pre-bone cartilage and stromal progenitor, sorted by CD200^−^CD51^+^Thy1^−^Ly51^−^CD105^−^, 1.2% ± 0.9%), PCPs (pro-chondrogenic progenitors, sorted by CD200^+^CD51^+^Thy1^+^Ly51^−^ CD105^+^, 2.5% ± 1.0%), BLSPs (B cell lymphocyte stromal progenitors, sorted by CD51^+^Thy1^+^Ly51^−^CD105^−^, 2.6% ± 1.5%), and THYs (CD200^−^CD51^+^Thy1^+^Ly51^−^CD105^+^, 0.4% ± 0.6%) (Figures 2G–2K). More importantly, the EYFP^+^CD200^+^CD51^+^Thy1^−^Ly51^−^CD105^−^CD45^−^ Ter119^−^CD31^−^ cohort from the *Col10a1^EYFP^* mice was fractionated by fluorescence-activated cell sorting (FACS) and transplanted under the kidney capsule of immunodeficient mice. These cells formed bony organoids with the properties of multipotential differentiation and supporting hematopoiesis (Figure S2F), indicating that HC descendants contained functional CD200^+^CD51^+^ SSCs. Taken together, HC-derived *Sstr2^+^*or *Pdgfrb^high^Kitl^low^* mpSSCs may represent a population of functional murine SSCs that specifically reside in the metaphysis.

### MpSSCs Are Bona Fide SSCs

We further investigated whether HC-derived *Pdgfrb^high^Kitl^low^* cells met the criteria for true murine SSCs. To specifically label the HC-derived *Pdgfrb*^+^ cells, we developed a sequential genetic lineage tracing system by generating *Col10a1-Cre;Pdgfrb^LSL-Dre^;ROSA26^LSL-EYFP/RSR-tdTomato^* (*CYPT*) mice, in which two recombination events occurred. All HC progenies were labeled with the Cre reporter *ROSA26^LSL-EYFP^,* which was triggered by *Col10a1-Cre*. Simultaneously, Cre excised the loxP-floxed STOP sequence before Dre at the *Pdgfrb^LSL-Dre^* locus. Thus, HC progeny could be recorded by the Dre reporter *ROSA26^RSR-tdTomato^* when they expressed *Pdgfrb* (Figure. 3A). Therefore, EYFP^+^tdTomato^+^ cells went through two stages, first expressing *Col10al* and then *Pdgfrb*, while EYFP^+^tdTomato^−^ offspring did not undergo *Pdgfrb* expression, even though they shared the same HC origin (Figure 3B). We verified no cross-reaction of Cre on Rox sites and efficient transcriptional blocking of the STOP sequence in *Col10a1-Cre;ROSA26^RSR-tdTomato^*(*CT*) and *Pdgfrb^LSL-Dre^;ROSA26^RSR-tdTomato^* (*PT*) mice, respectively (Figure S4A). Whole-mount fluorescent imaging showed that in the P1 *CYPT* rib, EYFP expression emerged at the HZ and continued throughout the trabecular zone, whereas tdTomato expression began at the top of the metaphysis and completely overlapped with EYFP expression (Figure S4B). Multicolor IF staining of the tibia showed that at P1, EYFP^+^tdTomato^+^ cells mainly resided in the metaphysis (Figure S4C).

**Figure 3.**
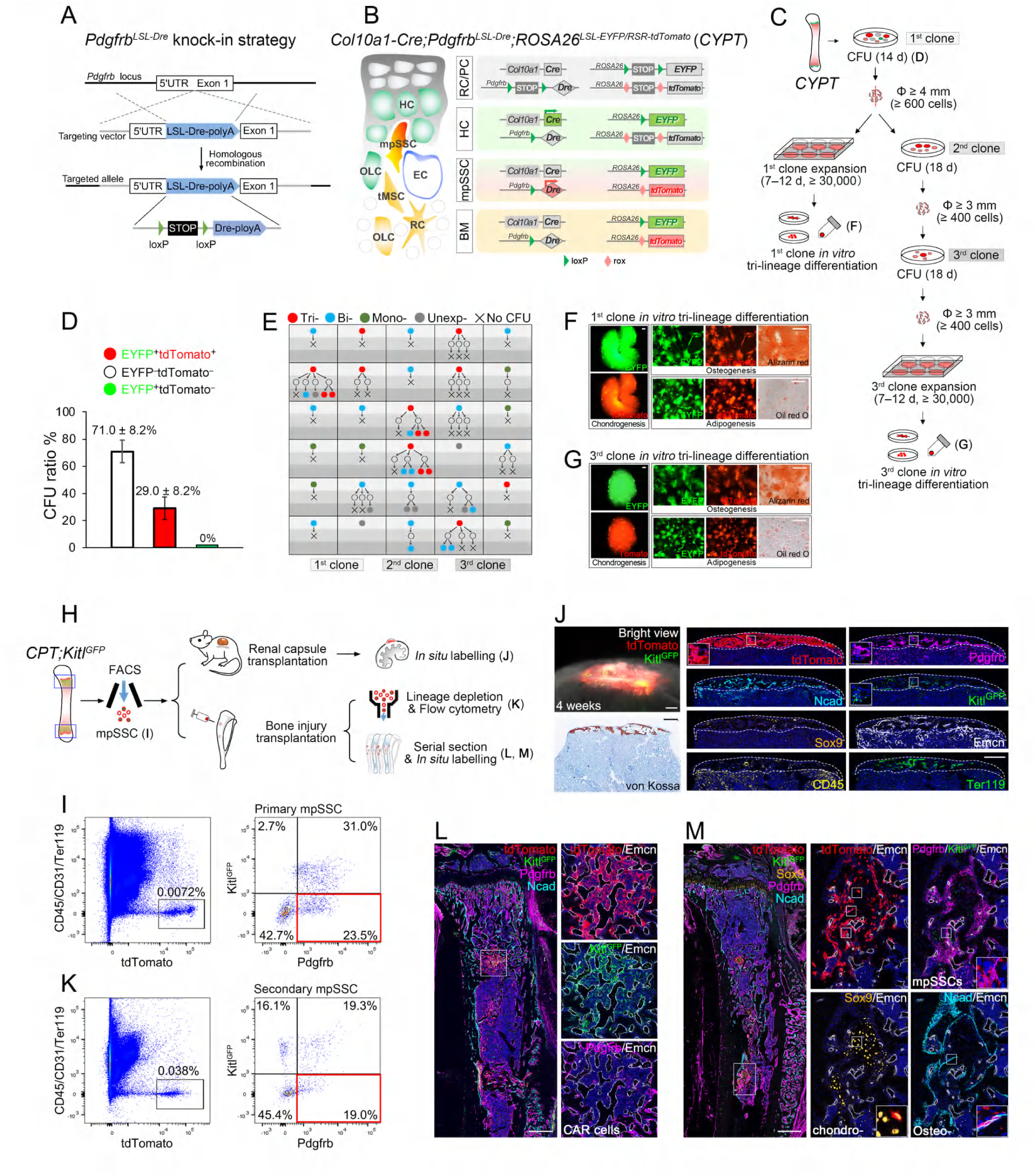
MpSSCs Are Bona Fide SSCs. (A) Strategy generating *Pdgfrb^LSL-Dre^* knock-in alleles by CRISPR-Cas9 technology. (B) Scheme showing the HC fate map. Within the growth plate of the *CYPT* mice, all HCs and their progeny are labeled by *Col10a1-Cre* triggered *ROSA26^LSL-EYFP^*. *Pdgfrb* is expressed when HC adopts the mpSSC fate, and *ROSA26^RSR-tdTomato^*expression marks Pdgfrb*^+^* mpSSC and their descendant cells. RC, resting chondrocyte; PC, proliferating chondrocyte; BM, bone marrow. (C) Scheme showing *in vitro* successional CFU formation and multilineage differentiation protocol for examining stem cell-like clones of P30 *CYPT* mice. (D) The ratio of 1^st^ CFUs (mean ± SD, 288 clones from 5 independent experiments). (E) Diagram showing the results of 30 clones in successional CFU formation and multilineage differentiation. Tri-, trilineage differentiation; Bi-, bilineage differentiation; Mono-, monolineage differentiation; Unexp-, failed population expansion to 30,000 cells. (F, G) Representative multilineage differentiation results of the 1^st^ generation clone and the 3^rd^ generation clone derived from a single stem cell-like EYFP^+^tdTomato^+^ cell. Scale bar, 100 μm. (H) Scheme showing the functional characterization of HC-derived Pdgfrb^high^Kitl^low^ mpSSCs by renal capsule transplantation model and bone injury transplantation model. Blue frames indicate the metaphyseal regions of P15–P30 *CPT;Kitl^GFP^* bones used for FACS. (I) The FACS plots shown are representative experimental results for sorting primary tdTomato^+^Pdgfrb^high^Kitl^low^ mpSSCs. (J) Representative wholemount fluorescent image, histology, and multicolor IF staining of bony organoids formed by tdTomato^+^Pdgfrb^high^Kitl^low^ mpSSCs 4 weeks after renal capsule transplantation. Dashed lines indicate the margins of tdTomato^+^ bony organoids. White arrows indicate a tdTomato^+^Pdgfrb^high^Kitl^low^ population. Scale bars, 200 μm. (K) The flow cytometry plots shown are representative experimental results for analyzing secondary tdTomato^+^Pdgfrb^high^Kitl^low^ mpSSCs extracted from callus 14 days after bone injury transplantation. (L, M) Representative multicolor IF staining of recipient tibias 14 days after tdTomato^+^Pdgfrb^high^Kitl^low^ mpSSCs transplantation. Serial sections cut at the callus formation site (white frames) show tdTomato^+^Pdgfrb^high^Kitl^low^ mpSSCs, tdTomato^+^Pdgfrb^+/–^Kitl^high^ CAR cells, tdTomato^+^Sox9^+^ chondrocytes (chondro-), and tdTomato^+^Ncad^+^ osteoblasts (osteo-) that are indicated by frames at high magnifications. Scale bars, 500 μm.

Next, we used an *in vitro* successional CFU and multilineage differentiation model to examine whether the HC-derived lineage contained stem cell-like cells (Figure 3C). The CFU-F assay of whole bone marrow cells isolated from *CYPT* mice revealed that EYFP^+^tdTomato^+^, but not EYFP^+^tdTomato^-^ cells, formed CFUs, suggesting that a history of *Pdgfrb* expression was required for HC progeny to acquire the capability of colony formation (Figure 3D). Furthermore, 9/30 clones exhibited trilineage differentiation potential. Even after two passage generations, trilineage differentiation potential was retained in 3/30 EYFP^+^tdTomato^+^ clones (Figures 3E–3G).

We further assessed the *ex vivo* differentiation and self-renewal ability of these tdTomato^+^Pdgfrb^high^Kitl^low^ cells by transplanting them under the renal capsule and into the bone injury sites (Figure 3H). TdTomato^+^Pdgfrb^high^Kitl^low^ mpSSCs were isolated from the metaphyseal bones of *Col10a1-Cre;Pdgfrb^LSL-Dre^;ROSA26^RSR-tdTomato^;Kitl^GFP^*(*CPT;Kitl^GFP^*) mice by FACS and transplanted under the kidney capsule of immunodeficient mice (Figure 3I). After 4 weeks, tdTomato^+^Pdgfrb^high^Kitl^low^ cells formed bony organoids, which were positive for von Kossa staining and contained Sox9^+^ chondrocytes, Ncad^+^ OLCs, Pdgfrb^+^ or Kitl^+^ stromal cells, host-derived blood vessels, and CD45^+^ or Ter119^+^ hematopoietic cells (Figure 3J). Notably, a small portion of tdTomato^+^ cells was Pdgfrb^high^Kitl^low^ (Figure 3J). This suggests these cells’ self-renewal ability, which was further validated in the bone injury transplantation model. The exogenous callus derived from the fractionated tdTomato^+^Pdgfrb^high^Kitl^low^ cells retained the tdTomato^+^Pdgfrb^high^Kitl^low^ mpSSC compartment (Figure 3K) and gave rise to the other three compartments, including Ncad^+^ OLCs, Sox9^+^ chondrocytes, and Pdgfrb^+/–^ Kitl^high^ stromal cells (Figures 3L and 3M). In contrast, tdTomato^+^Pdgfrb^+^Kitl^+^ cells, the putative immediately downstream descendants of Pdgfrb^high^Kitl^low^ cells, failed to repopulate their own compartments in the grafted bones, possibly due to defects in cell proliferation or survival (Figure S4E). In summary, the above results indicate that HC- derived Pdgfrb^high^Kitl^low^ cells possess clonal multipotency and self-renewal capability, and are true functional SSCs.

### MpSSCs Generate Most HC Descendants

Given the inherent nature of SSCs, we hypothesized that mpSSCs are likely to produce the majority of HC progeny. We mapped all HC descendants in *CPYT* mice at different time points in detail. At P1, the ratio of EYFP^+^tdTomato^+^ cells to EYFP^+^tdTomato^−^ cells was 1:2.19 (n = 6, Figure 4A). However, the ratio of EYFP^+^tdTomato^+^ cells to EYFP^+^tdTomato^−^ quickly reversed at P7 as shown by EYFP^+^tdTomato^+^ cells accounting for most of total EYFP^+^ HC-descendants (77.1% ± 1.7%, n = 6) at multiple sites (Figure. 4A), such as 57.4% ± 5.1% in cortical endosteum, 80.0% ± 1.4% in trabeculae, and 97.7% ± 1.7% in bone marrow (n = 7) (Figures 4B and 4C). The number of EYFP^+^tdTomato^+^ cells continued to increase and peaked at P30 (Figures 4A and S5A). Although both the number of EYFP^+^tdTomato^+^ and EYFP^+^tdTomato^−^ cells gradually decreased along with the cessation of endochondral ossification at P180 as well as aging, the percentage of EYFP^+^tdTomato^+^ cells was extremely high over time (91.1% ± 1.8%, 96.7 ± 0.6%, 98.5% ± 0.6%, and 98.8% ± 0.6% at P15, P30, P180, and P480, respectively, n = 6, Figures 4A and S5B).

**Figure 4.**
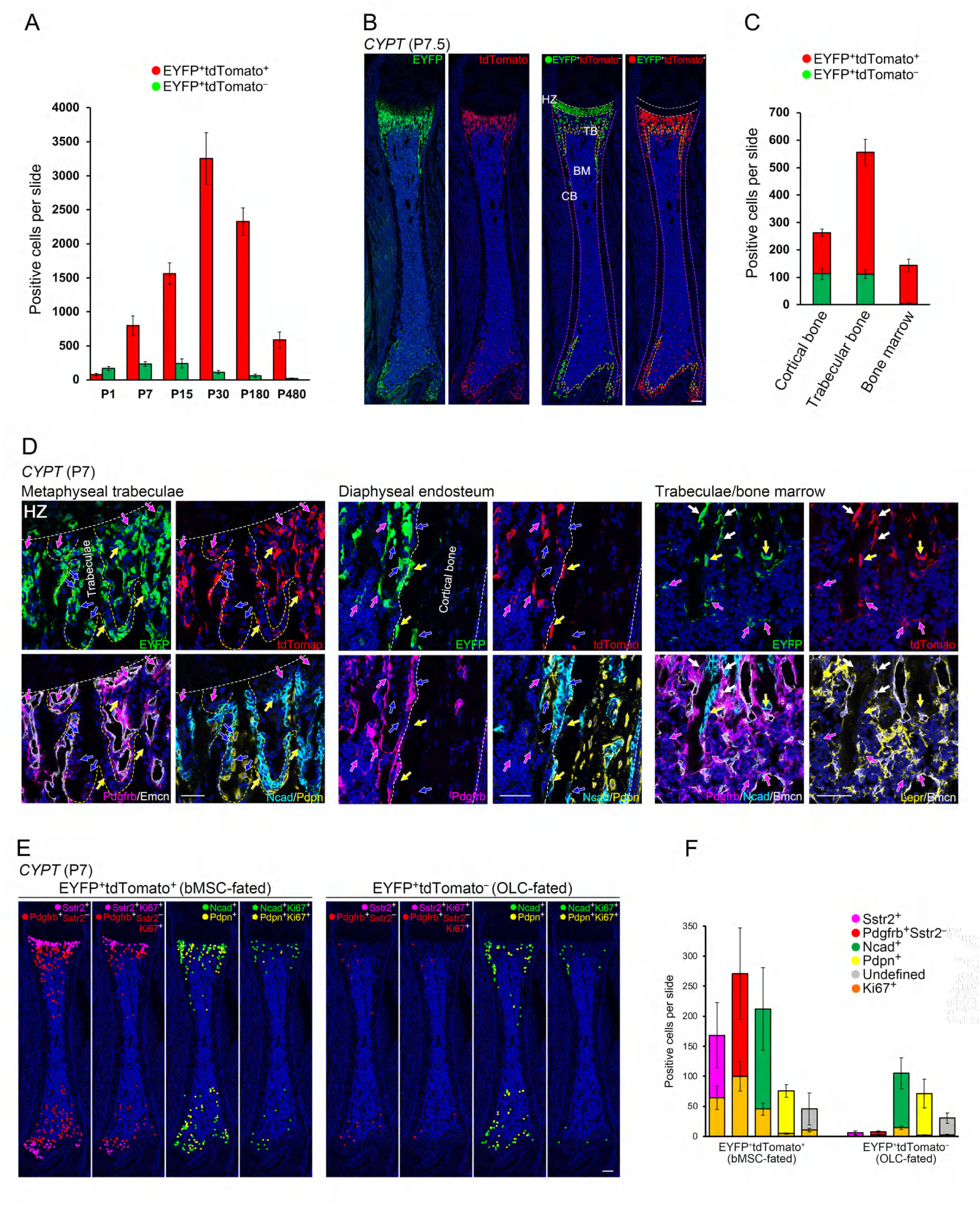
MpSSCs Generate Most HC Descendants. (A) Quantification of EYFP^+^tdTomato^+^ and EYFP^+^tdTomato^−^ cells in femur sections of *CYPT* mice at different ages. Data are presented as mean ± SD per slide (n = 6). (B) Mapping of EYFP^+^tdTomato^+^ and EYFP^+^tdTomato^−^ cells in femur sections of P7.5 *CYPT* mice. White, magenta, and yellow dashed lines outline the hypertrophic zone, cortical bone, and trabecular bone, respectively. Red dot-represented EYFP^+^tdTomato^+^ cells and green dot-represented EYFP^+^tdTomato^−^ cells are superimposed on the same P7.5 *CYPT* femur section. HZ, hypertrophic zone; TB, trabecular bone; CB, cortical bone; BM, bone marrow. Scale bar, 200 μm. (C) Quantification of EYFP^+^tdTomato^+^ and EYFP^+^tdTomato^−^ cells within the cortical bone, trabecular bone, and bone marrow as shown in (B). Data are presented as mean ± SD per slide (n = 7). (D) Mapping of two-fated HC descendants in femur sections of P7 *CYPT* mice. Dashed lines outline the chondro-osseous junction, a metaphyseal trabecula, and the cortical bone. Magenta arrows, EYFP^+^tdTomato^+^Pdgfrb^+^ cells; yellow arrows, EYFP^+^tdTomato^+^Ncad^+^/Pdpn^+^ cells; blue arrows, EYFP^+^tdTomato^−^Ncad^+^/Pdpn^+^ cells; white arrows, EYFP^+^tdTomato^+^Pdgfrb^+^Lepr^−^ cells. Scale bar, 50 μm. (E) Color dot-represented cell compartments are superimposed on femur sections of the P7 *CYPT* mice. Colored pentagons represent Ki67-positive cells from the same cell compartment. Scale bar, 200 μm. (F) Quantification of each cell compartment as shown in (E). Data are presented as mean ± SD per slide (n = 4). See also Figure S3.

In accordance with the multipotential ability of mpSSCs, the IF analysis of P7 *CPYT* femurs revealed that EYFP^+^tdTomato^+^ cells accounted for almost all bMSC lineages, including Sstr2^+^ mpSSCs and their downstream Sstr2^−^Pdgfrb^+^ or Pdgfrb^+^Lepr^+/–^ bMSCs in the metaphysis and bone marrow, as well as a large part of Ncad^+^ or Pdpn^+^ OLCs (Figures 4D and 4E). In contrast, the EYFP^+^tdTomato^−^ population was largely composed of Ncad^+^ or Pdpn^+^ OLCs, similar to the findings at P1 (Figures 4D–4F and S4D). Furthermore, Ki67^+^ proliferative cells were highly enriched in EYFP^+^tdTomato^+^ populations, including Sstr2^+^, Sstr2^−^Pdgfrb^+^, and Ncad^+^ cells (Figures 4E and 4F). This could partially contribute to the robust increase in EYFP^+^tdTomato^+^ cells. Additionally, a few EYFP^+^tdTomato^+^ adipocytes were observed in the bone marrow of the P180 *CYPT* mice (Figure S5C). Notably, EYFP^+^tdTomato^+^Sstr2^+^ cells remained at P180, reflecting their long-term survival or self-renewal ability (Figure S5B).

Taken together, HC-derived mpSSCs survive long-term and give rise to most HC descendant cells, including bMSCs and OLCs.

### Hgs Deletion in HCs Impairs the Conversion of HCs to MpSSCs

The abrupt changes in cellular morphology and transcriptional signatures from HCs to mpSSCs raised the question of how HC acquired mpSSC identity. Next, we used Gene Ontology (GO) and Kyoto Encyclopedia of Genes and Genomes (KEGG) analyses to investigate the biological processes, cellular components, and signaling pathways enriched in HCs, particularly HC-1, which seemed to be responsible for initiating the HC to mpSSC conversion (Figures 5A–5C). We focused on endosomal transport pathways that were enriched in GO terms (biological processes and cellular components) and KEGG analyses, as the endosomal transport pathway is essential for diverse receptor-mediated signal transduction and is widely implicated in various cellular processes (Szymanska et al., 2018) (Figures 5A–5C). We found that Hgs, a pivotal component of the endosomal sorting complex, was enriched in the HC-1 subset and involved in multiple cellular transport biological processes (Figure 5D). Consistently, the IF analysis showed that Hgs largely overlapped with the HC-1 marker Tnxip, but less with the HC-2 marker F13a1 or the HC-3 marker Pla2g10 in the femurs of *Col10a1-Cre;Hgs^+/+^* mice (Figure 5E). To evaluate the role of Hgs in regulating the fate of HCs, we generated *Col10a1-Cre;Hgs^fl/fl^* mice to delete the *Hgs* gene in HCs. At P7, *Hgs* mutant femurs displayed an irregular arrangement of HCs, with a marked decrease in Txnip^+^ HCs and an increase in F13a1^+^ or Pla2g10^+^ HCs (Figure 5E), indicating a skewed cell population distribution within the HCs.

**Figure 5.**
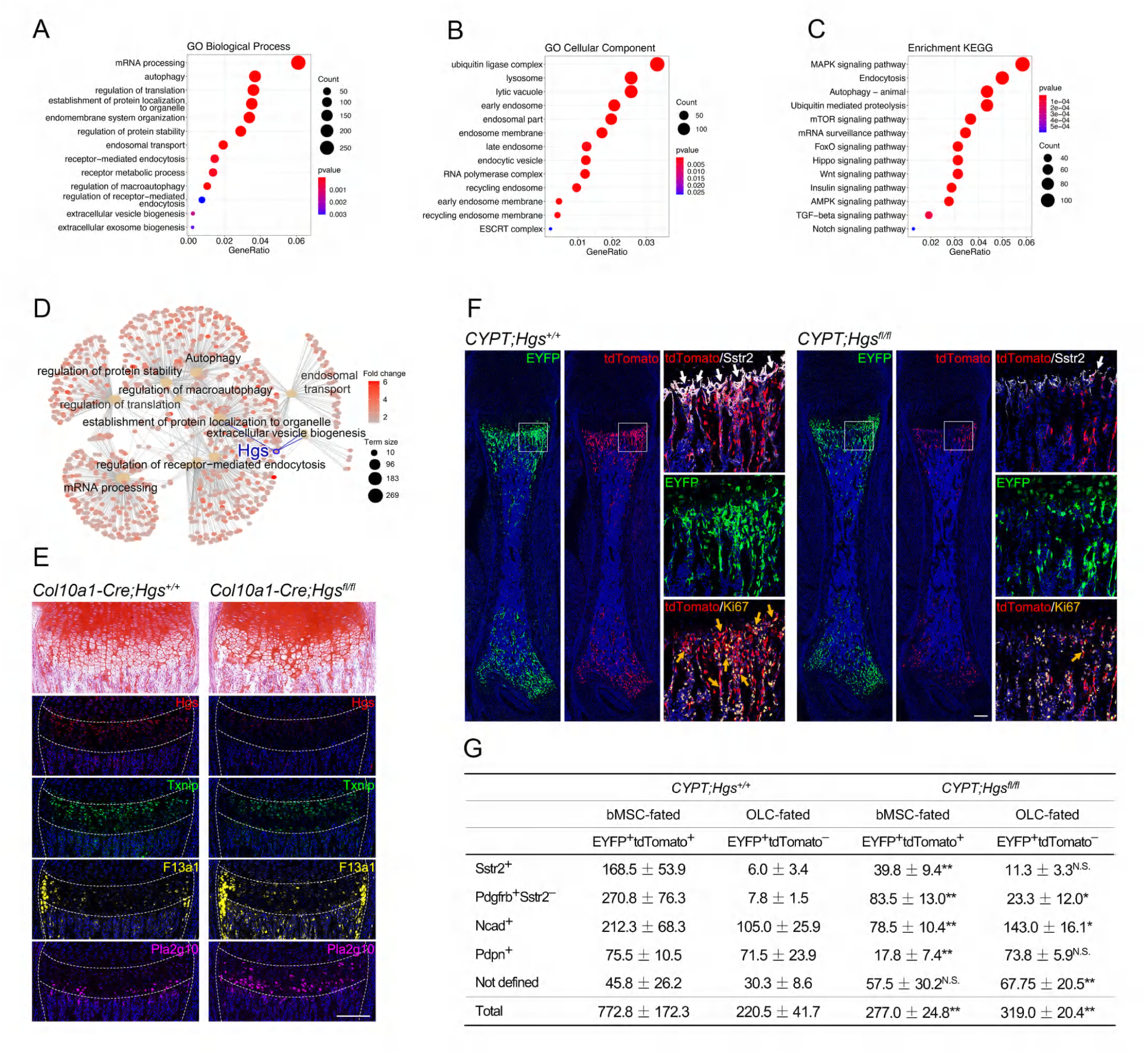
Hgs Deletion in HCs Impairs the Conversion of HCs to MpSSCs. (A-D) Single-cell analysis reveals that Hgs is important in HC-1 cells. Dotplot (A-C) and cnetplot (D) of significant biological pathways enriched by high-expressed genes of HC-1 cells. The Hgs-related biological processes (A, D), cellular component (B), and KEGG pathways (C) are enriched in HC-1 cells. The *Hgs* gene and its related biological processes are highlighted in blue in cnetplot (D). (E) Representative safranine O staining and multicolor IF staining indicating altered HC arrangement and HC subpopulations in femurs of P7 *Col10a1-Cre;Hgs^fl/fl^* mice. Scale bar, 200 μm. (F) Representative multicolor IF staining of femurs indicating impaired conversion of HCs to mpSSCs in P7 *CYPT;Hgs^fl/fl^*mice. White arrows indicate tdTomato^+^Sstr2^+^ mpSSCs, gold arrows indicate tdTomato^+^Ki67^+^ proliferative cells. Scale bars, 200 μm. (G) Quantification of two-fated HC descendants in P7 femurs of *CYPT;Hgs^+/+^* and *CYPT;Hgs^fl/fl^* mice. Data are presented as mean ± SD per slide (n = 4). *p < 0.05, **p < 0.01, N.S., no significance.

Next, we examined the effect of Hgs deletion on the conversion of HCs to mpSSCs using *Col10a1-Cre;Pdgfrb^LSL-Dre^;ROSA26^LSL-EYFP/RSR-tdTomato^;Hgs^fl/fl^* (*CYPT;Hgs^fl/fl^*) mice. EYFP^+^tdTomato^+^Sstr2^+^ mpSSCs in *Hgs* knockout mice decreased by 76.4% at P7 (Figures 5F and 5G). Furthermore, the number of total EYFP^+^tdTomato^+^ populations, including Sstr2^−^Pdgfrb^+^ bMSC and Ncad^+^ or Pdpn^+^ OLCs, decreased accordingly (Figure 5G). These data demonstrated that Hgs is essential for the conversion of HCs to mpSSCs. Interestingly, the population of EYFP^+^tdTomato^−^Ncad^+^ OLCs, which was directly transdifferentiated from HCs, increased by 1.36 fold in *CYPT;Hgs^fl/fl^*mice (Figure 5G). However, the increase in EYFP^+^tdTomato^−^ OLCs was insufficient to rescue the decrease in the overall pool size of EYFP^+^ HC descendants (Figure 5G), indicating the key role of mpSSCs in generating HC progeny.

### MpSSC Loss Compromises Trabecular Bone Formation and Bone Regeneration

Taking advantage of the pronounced loss of mpSSCs caused by HC-specific *Hgs* deletion, we assessed the effect of mpSSC loss on bone formation. At P30, the *Hgs* knockout mice displayed virtually no bMSCs and a remarkable decrease in the OLC population (Figures 6A and 6B). Micro-computed tomography demonstrated a defect in trabecular bone formation in Hgs-knockout mice, as evidenced by a lower bone marrow density (BMD) and bone mass (BV/TV) in Hgs-deficient trabecular bone than in control mice (Figures 6C and 6D). Consistently, Hgs-deficient mice displayed a significant decrease in the trabecular number (Tb. n) and thickness (Tb. th), resulting in increased trabecular spacing (Tb. sp) (Figures 6C and 6D). We also found that these HC-derived bMSCs constitute a reservoir of putative SSCs that robustly contribute to bone regeneration, as shown by the emergence of injury-associated EYFP^+^Pdgfrb^+^CD200^+^CD51^+^CD105^−^ SSCs at the metaphyseal and diaphyseal injury sites in the drill hole model (Figures 6E and 6F). However, in *Hgs* knockout mice, the loss of HC-derived Pdgfrb^+^ bMSCs or CD200^+^CD51^+^ SSCs compromised injury-associated osteogenesis and bone callus formation (Figures 6F and 6G). Collectively, these results suggest that mpSSCs play essential roles in trabecular bone formation and bone regeneration by supplying bMSCs and OLCs.

**Figure 6.**
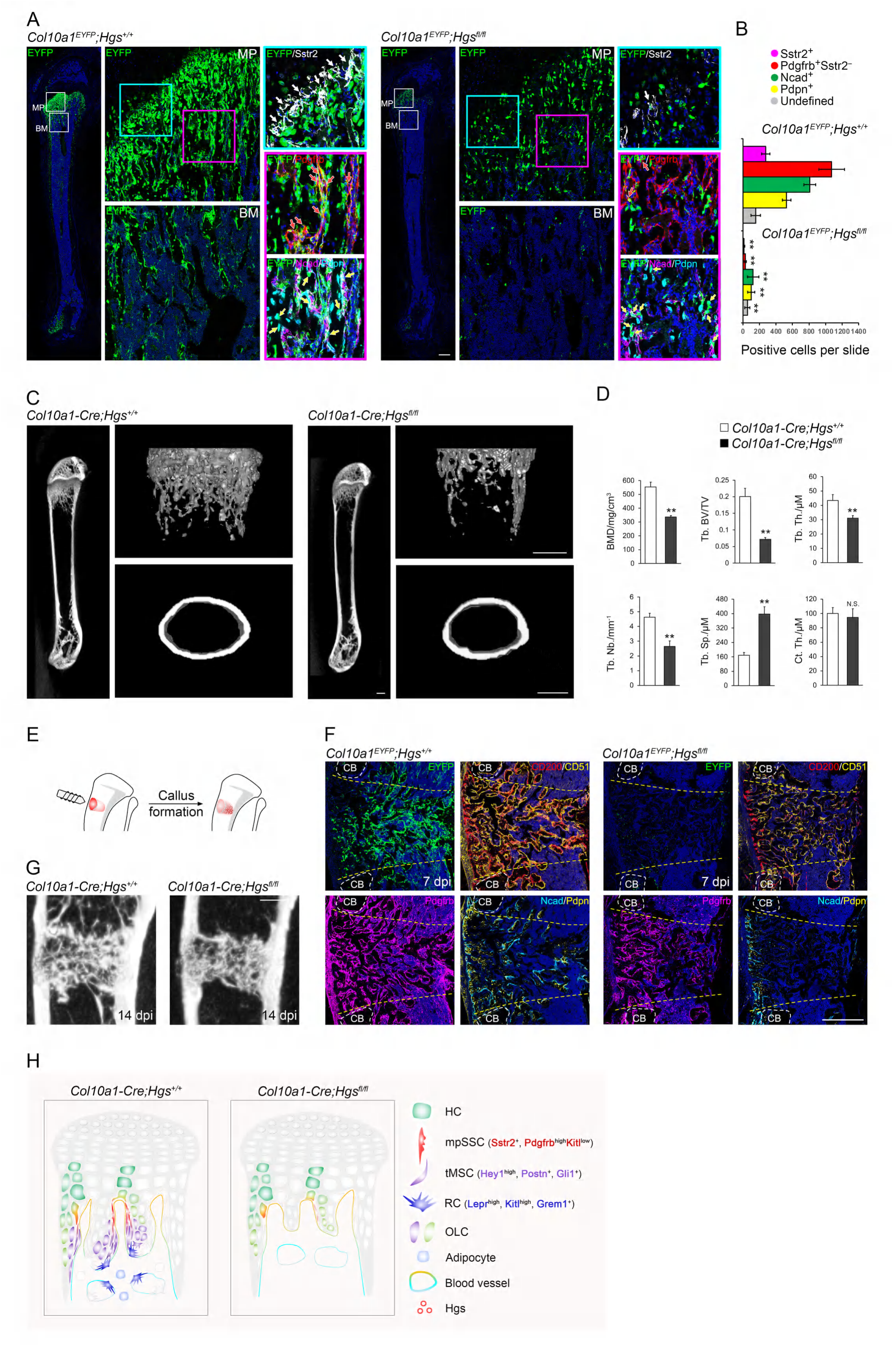
MpSSC Loss Compromises Trabecular Bone Formation and Bone Regeneration. (A) Representative multicolor IF staining showing significantly decreased HC-derived mpSSCs, bMSCs, and OLCs in femurs of 1-month-old *Hgs^fl/fl^* mice. The framed areas are shown adjacently at higher magnifications. MP, metaphysis; BM, bone marrow. (B) Quantification of each cell compartment as shown in (A). Data are presented as mean ± SD per slide (n = 6). **p < 0.01. (C) Representative reconstructed μCT images of the femur showing impaired trabecular bone formation in 1-month-old *Col10a1-Cre;Hgs^fl/fl^* mice. (D) Quantification of metaphyseal parameters of the femur as shown in (C). Data are presented as mean ± SD (n = 6). BMD, bone mineral density; Tb. BV/TV, trabecular bone surface area/bone volume; Tb. Th., trabecular thickness; Tb. Sp., trabecular spacing; Tb. Nb., trabecular number; Ct.Th., cortical thickness. **p < 0.01, N.S., no significance. (E) Scheme showing the bone drilling surgery that induces intra-marrow callus formation. (F) Representative multicolor IF staining showing decreased CD200^+^CD51^+^, Pdgfrb^+^, and Ncad^+^/Pdpn^+^ cells within the intra-marrow calluses of *Col10a1^EYFP^*;*Hgs^fl/fl^* tibia on day 7 post-injury (7 dpi). White dashed lines outline the cortical bone where the bone drilling surgery was performed. Yellow dashed lines outline the intra-marrow calluses. (G) Representative reconstructed μCT images of calluses showing impaired ossification of intra-marrow calluses in 1.5-month-old *Col10a1^EYFP^*;*Hgs^fl/fl^* tibia on day 14 post-injury (14 dpi). Scale bar, 500 μm. (H) Scheme summarizing the findings of this study.

## DISCUSSION

Here, we identified a previously unappreciated mpSSC population, which were derived from HCs and involved in trabecular bone formation and regeneration, establishing a new cellular paradigm for trabecular bone formation (Figure 6H).

A prominent character of mpSSCs is their exclusive cellular origin. Theoretically, marrow SSCs are thought to be derived from perichondral cells that migrate into marrow and HCs that de-differentiate at the perinatal stage (Ambrosi et al., 2019; Serowoky et al., 2020). Our lineage tracing results demonstrated that all mpSSCs are the direct descendants of HCs, suggesting that the de-differentiation of HC into mpSSC is a key process bridging chondrogenesis and osteogenesis during endochondral ossification. MpSSCs emerged at perinatal stage, peaked at the age of 1 month, and declined after the age of 3 months. The temporal and spatial pattern of mpSSCs corresponds to the active HC transition and robust trabecular bone formation in the postnatal stage. Thus, adopting a mpSSC fate not only endows HCs with greater differentiation potential but also enables HCs to survive longer and produce progeny that participate in postnatal bone formation. Consistently, the physiological function of mpSSC is critical for trabecular bone formation, as a reduced mpSSC pool caused by loss of Hgs led to significantly compromised trabecular bone formation. Furthermore, we revealed that HC-derived mpSSCs stand at the pinnacle of the HC dedifferentiation trajectory. Previous lineage tracing studies have shown that HCs can give rise to over half of all Lepr^+^ bone marrow stromal cells, as well as marrow-associated skeletal stem and progenitor cells (SSPCs) marked by Pdgfra and Lepr (Long et al., 2022; Shu et al., 2021). However, these two populations mainly reside within diaphyseal bone marrow and it remains undetermined whether they are bona fide SSCs due to the lack of the *in vitro* and *in vivo* functional assessment. When comparing to the localization and transcriptional profiling of HC progeny defined here, both HC-derived Lepr^+^ bone marrow stromal cells and SSPCs are possibly downstream offspring of Pdgfrb^high^Kitl^low^ mpSSCs.

As the first formally defined functional SSC population in the metaphyseal trabeculae, mpSSCs serve as a reservoir for previously reported metaphyseal mesenchymal progenitors and diaphyseal bMSCs. In the metaphysis region, Grem1^+^, Gli1^+^, Osterix^+^, and Pdgfra^+^Pdgfrb^+^Hey1^+^ label multiple types of metaphyseal mesenchymal progenitors that contribute to trabecular osteoblasts and bone marrow stromal cells (Mizoguchi et al., 2014; Shi et al., 2017; Sivaraj et al., 2021; Worthley et al., 2015). Additionally, the lower regions of the metaphysis and diaphyseal marrow harbor several overlapping bone marrow stromal cell populations marked by Lepr, Kitl, and Cxcl12 (Ding et al., 2012; Sugiyama et al., 2006; Zhou et al., 2014a). We found that mpSSCs expressed high levels of Pdgfrb and Hey1, but little or very low levels of Kitl, Lepr, Grem1, Sp7/Osterix, and Gli1, whereas tMSCs and RCs, two mpSSCs downstream populations, expressed Postn, Gli1, Sp7, Lepr, Cxcl12, Kitl, and Grem1 to varying extents. We propose that HC-derived Pdgfrb^high^Kitl^low^ mpSSCs represent a subpopulation of Hey1^+^Pdgfrb^+^ bMSCs and could give rise to Gli1^+^, Sp7^+^, Lepr^+^, Cxcl12^+^, and Grem1^+^ progenitor cells, therefore serving as a key cellular source for skeletal progenitors and bone marrow stromal cells. Notably, mpSSCs give rise to a fraction but not all of the aforementioned bMSCs populations as evidenced by sc-RNA-seq, multiple IF staining and CFU-F results. This raised a possibility of whether other SSC populations with a non-HC origin exist in the lower metaphyseal and diaphyseal bone marrow. A candidate SSC population may reside at the cortical bone endosteum, where CD200^+^CD51^+^CD90^−^Ly51^−^CD105^−^ cells are also enriched shown by multiple IF analysis. On the other hand, other SSC populations may not be necessary, as the extraordinary cellular plasticity of bMSCs enable them to reacquire the ability of SSCs as shown in bone regeneration (Knopf et al., 2011; Storer et al., 2020).

The nature of HC-derived mpSSC provides insight into the heterogeneity and interactions of diverse previously reported SSCs at different sites. Previous studies have shown that epiphyseal growth plate houses the PTHrP^+^ SSCs that reside at resting zone and give rise to HCs, bone marrow stromal cells, and osteoblasts (Mizuhashi et al., 2018; Newton et al., 2019). Notably, only a few offspring of PTHrP^+^ SSCs can extend into the metaphysis beyond the age of 2 months (Mizuhashi et al., 2018), whereas the active HC-to-mpSSC transition mainly occurs before P30, suggesting that Sstr2^+^ mpSSC are not the descendants of PTHrP^+^ SSCs. Together with the fact that the offspring of periosteal SSCs are confined to periosteum, it seems that SSCs at distinct sites exert their distinct functions in a site-specific manner. However, a recent study showed that periosteal SSCs control the function of the growth plate resting PTHrP^+^ SSCs by periosteal SSCs-derived Indian hedgehog (Tsukasaki et al., 2022). Therefore, although the cellular crosstalk between these SSCs is still lack of evidence, the functional interaction between mpSSCs and other previously identified SSCs need to be clarified in future studies. Nevertheless, taken together with murine SSCs identified in the growth plate, periosteum, and articular surface (Debnath et al., 2018; Mizuhashi et al., 2018; Murphy et al., 2020; Pineault et al., 2019), these findings support that multiple SSCs, which have distinct temporal and spatial contexts and origins as well as different functional properties, work in concordance to achieve longitudinal bone development.

## ACKNOWLEDGMENTS

This work was supported by the National Natural Science Foundation of China (31630093 to X.Y., 31871476, 31571512 to G.Y.), Beijing Nova Program (Z161100004916146 to G.Y.).

## AUTHOR CONTRIBUTIONS

X.Y., Y.T., and G.Y. conceived and designed the project. G.Y., Q.H., X.G., J.L., R.L., L.Z., W.T., W.L., H.L., M.T., Y.Q., Y.P., L.D., K.S., Z.X., C.L. and M.L. performed experiments. X.Y., Y.T., and G.Y wrote the manuscript with significant input from B.Z.. G.Y., Q.H. and X.G. contributed equally to the study.

## DECLARATION OF INTERESTS

The authors declare no competing interest.

## EXPERIMENTAL MODEL AND SUBJECT DETAILS

### Mice

The lines used in this study were as follows: *Col10a1^int2^-Cre* (*Col10a1-Cre*), *ROSA26^LSL-EYFP^*, *ROSA26^LSL-Confetti^*, *ROSA26^LSL-ZsGreen^*, *ROSA26^LSL-tdTomato^ ROSA26^RSRtdTomato^*, *Kitl^GFP^*, and *Ihh-mKate2^tomm20^-Dre* (Ding et al., 2012; Li et al., 2022; Madisen et al., 2010; Wang et al., 2017; Yang et al., 2014a). For the renal capsule transplantation experiment, 6–8-week-old NOD SCID immunodeficient mice were used as recipient hosts.

We generated a Pdgfrb-LoxP-STOP-LoxP-Dre knock-in line by homologous recombination with CRISPR-Cas9. A LoxP-5×PloyA-LoxP-Dre cassette was inserted after the translational start codon of the *Pdgfrb* gene. F0 adults were outcrossed to the wild type, and F1 embryos were genotyped to assess germline transmission in the presence of insert fragments.

All experiments were performed according to the guidelines of the Laboratory Animal Center of AMMS (approval number: IACUC-DWZX-2020-001).

### Cell Preparation for Single Cell RNA Sequencing

P7.5 *Col10a1^ZsGreen^* mice were used for single-cell analysis of HCs and their descendants. The femur and tibia were dissected and cleaned by brief digestion and brushing in 0.2% collagenase type I (GIBCO, 17100017) for 30 min. To exclude cell interference from the secondary ossification center and other sites of the growth plate, we only collected the proximal part of the femur and distal part of the tibia, with half of the growth plate cartilage excised. To help release the embedded cells, cortical bone, trabecular bone, and hypertrophic cartilage were stripped and sliced into pieces using a syringe needle under a stereoscope. The samples were then digested in 0.2% collagenase type I and 0.1% collagenase type II (GIBCO, 17100015) for 2–3 h at 37 °C under gentle agitation. The liberated cells were pipetted, filtered, neutralized, and centrifuged every 20 min. Dead cells were removed using a kit (130090101, STEMCELL Technologies). The collected cells were prepared to establish two types of single-cell transcriptomes. The first was the unsorted scRNA-seq data, which were constitutive from skeletal cells of the whole bone with hematopoietic and endothelial cells briefly depleted by CD31/CD45/Ter119-coated magnetic beads (Biotin anti-mouse CD31 Antibody, BioLegend, 102503; Biotin anti-mouse CD45 Antibody, Biolegend, 103104; Biotin anti-mouse TER-119 Antibody, Biolegend, 116203; EasySep™ Mouse Streptavidin RapidSpheres™ Isolation Kit, STEMCELL Technologies, 19860). The other was composed of ZsGreen-positive cells sorted by FACS Aria III (BD Bioscience). Single cells were encapsulated in emulsion droplets using a chromium controller (10× Genomics). The scRNA-seq libraries were prepared using Chromium Single Cell 3ʹ Reagent Kits v3 (10× Genomics), according to the manufacturer’s protocol. The scRNA-seq libraries were evaluated and quantified using an Agilent 2100 Bioanalyzer/Fragment Analyzer 5300 and Qubit HS. The libraries were sequenced on a NovaSeq platform (Illumina) to generate 150 bp paired-end reads, according to the manufacturer’s instructions.

### Processing and Analysis of Single-cell RNA Sequencing Data

Raw data from five samples based on 10× Genomics were processed using the Cell Ranger software suite (v3.1.0) with reference genome mm10, and doublets were removed using Scrublet (v0.2.1) (Wolock et al., 2019). Hematopoietic and endothelial cells were removed from ZsGreen-sorted and unsorted scRNA-seq data. Cells with gene number more than 2000 or a percentage of mitochondrial genes lower than 15% were remained and projected into two dimensions by uniform manifold approximation and projection (UMAP) and clustered by K-means algorithm using Scanpy (Wolf et al., 2018) (v1.7.2) in the Python environment (v3.9). ZsGreen-expressing cells (UMI > 0) were extracted from the unsorted scRNA-seq data and integrated with ZsGreen positive cells sorted by FACS. The batch effect was removed using the cellranger aggr function. Based on the integrated ZsGreen+ data, cells expressing gene numbers less than 700 or a percentage of mitochondrial genes higher than 15% were filtered out. The remaining cells were projected into two dimensions by uniform manifold approximation and projection (UMAP) using a cellranger and clustered by the Leiden algorithm (resolution = 0.3) using Scanpy (Wolf et al., 2018) (v1.7.2) in the Python environment (v3.9). Slingshot trajectory inference was based on UMAP reduction dimensions. The Wilcoxon rank-sum test was performed to identify differentially expressed genes (DEGs) in each cell cluster, and the p-values were adjusted using the Benjamini-Hochberg correction. Genes with an adjusted p-value < 0.05, log (fold change) > 0.5, and expressed in over 30% of the cells in the cluster were selected for GO and KEGG enrichment analyses using the R package ClusterProfiler (v3.14.3) (Wu et al., 2021). Finally, signaling entropy was computed using the R package SCENT (v1.0.2) to infer differentiation potency at the single-cell level (Teschendorff and Enver, 2017).

### Micro-CT analysis

Femurs and tibiae were freshly dissected, fixed in 4% PFA, scanned, and analyzed using a micro-CT Skyscan 1076 μCT scanner (Bruker Corporation) and software. The scanner was set at a voltage of 80 kV, a current of 500 mA, and a resolution of 9.2 μm/pixel. Metaphyseal parameters of the femur, including bone mineral density (BMD), trabecular bone surface area/bone volume (Tb. BV/TV), trabecular thickness (Tb. Th.), trabecular spacing (Tb. Sp.), and trabecular number (Tb. Nb.), were measured from 0.5 mm to 2.3 mm below the lowest part of the metaphyseal growth plate. The cortical bone thickness (Ct. Th.) was measured from 2 mm to 2.5 mm below the lowest part of the metaphyseal growth plate of the femur. Longitudinal slices of the femurs and calluses of the tibiae were reconstructed at a thickness of 200 μm.

### Marrow Stromal Cell Isolation, Successional Clonal Formation, and Multilineage Differentiation

Bone marrow cells of P30 *Col10a1-Cre;Pdgfrb^LSL-Dre^;ROSA26^LSL-EYFP/RSR-tdTomato^* mouse tibia and femur were repeatedly flushed out using a 1-mL syringe. To help release the stromal cells residing between the metaphysical trabeculae, the growth plate cartilage was peeled off, and the metaphysis part of bones was cut off, gently disrupted using a mortar and pestle, and digested in 0.2% collagenase type I for 2–4 h. The liberated primary bone marrow and stromal cells were filtered, collected, isolated, and plated at a density of 2 × 10^5^ cells/10 cm dish to allow the formation of individual colonies after growth in DMEM (GIBCO, 11885084) containing 12% FBS (GIBCO, 12664025) and 1% antibiotic-antimycotic (GIBCO, 15240-062) at 37 °C under hypoxic conditions (5% O_2_, 5% CO_2_) for 2 weeks. Single clones with ≥600 cells (Ø ≥ 4 mm) were selected and labeled as 1^st^ clones. They were equally divided into two parts: one was re-plated in a 10 cm dish to allow the formation of 2^nd^ individual colonies, and the other part was expanded in 6-well plates to reach the cell population (≥30,000 cells) for further multi-lineage differentiation potential test of the 1^st^ clones. We generated 3^rd^ individual colonies (≥400 cells/Ø ≥ 3 mm) from 2^nd^ colonies (≥400 cells/Ø ≥ 3 mm) via the same re-plating protocol. For *in vitro* multi-lineage differentiation potential test, each expanded 1^st^ and 3^rd^ clone was split for differentiation into chondrocytes, osteoblasts, and adipocytes using a differentiation kit (STEMCELL Technologies, 05507, 05455, and 05504). The *in vitro* differentiation reported in this study is clonal. Images were taken using a Nikon ECLIPSE Ti-U inverted microscope system.

### Cell Isolation, FACS, and Renal Capsule Transplantation

Limb bones were dissected from mice at the indicated age. They were briefly digested with 0.2% collagenase type I at 37 °C for 30 min. After the removal of the muscle and ligament, the tibiae and femurs were dissociated by mechanical and enzymatic dissociation according to a published protocol (Gulati et al., 2018). Flow cytometry was performed using FACS Aria III. The total dissociated cells were blocked with rat IgG and stained with fluorochrome-conjugated antibodies against CD45 (BioLegend, 103154, 1:400), CD31 (BioLegend, 102528, 1:400), Ter119 (BioLegend, 116223, 1:200), CD140b (Biolegend, 136010, 1:400), CD51 (Biolegend, 104105, 1:200), CD200 (Biolegend, 123809, 1:400), CD105 (Biosciences, 740425, 1:200), Thy1.2 (Biolegend, 105335, 1:200), and Ly51 (Biolegend, 108313, 1:200) for fractionation by FACS. Dead cells were excluded using 7-AAD staining (Wolock et al., 2019). 200–400 murine SSCs from femurs and tibiae were enriched per P7–P10 *Col10a1^EYFP^*mouse. The 300–500 tdTomato^+^Pdgfrb^high^Kitl^low^ mpSSCs could be enriched from the fore/hind limb bones and scapulae per P10–P20 *CPT;Kitl^GFP^* mice. Sorted cells (6,000–10,000) were pelleted, resuspended in 2–3 μL of Matrigel (Corning, 356234), and then injected underneath the renal capsules of 6–8-week-old NOD SCID immunodeficient mice. After 4–6 weeks, the recipient kidneys were dissected for histological and immunohistochemical analysis.

### Bone Injury and Transplantation

To evaluate the contribution of bone marrow *mesenchymal stromal cells* to bone regeneration, a bone injury model was created by drilling a Ø 0.8 mm hole on the medial surface of the tibial shaft 2 mm beneath the growth plate. Compared with the fracture model, drilling holes at a relatively thin bone site can minimize the formation of cartilaginous calli, thus avoiding the influence of newly generated HC descendants via endochondral ossification. To transplant isolated cells to the bone injury site, 6–8-week-old NOD SCID immunodeficient mice were used as recipient hosts. First, a 26-gauge syringe needle was placed parallel to the tissue between the cortical bone and periosteum/fascia, making a hole at the metaphyseal medial tibial shaft 1 mm beneath the growth plate, which was covered by the periosteum/fascia. A 2^nd^ Ø 0.5 mm hole passing through the cortical bone and periosteum/fascia was then made by drilling at the diaphyseal medial tibial shaft, 3 mm beneath the growth plate. Using a 31-gauge insulin pen needle and a microinjector, 6,000–10,000 sorted cells resuspended in 2–4 μL Matrigel were injected into the bone marrow through the metaphyseal hole. After 10–14 days, recipient tibiae were dissected for flow cytometry or immunohistochemical studies. Before flow cytometry, non-skeletal cells were briefly depleted by CD31/CD45/Ter119-coated magnetic beads.

### Multiplex Immunofluorescent Assay

Samples of the hindlimb, rib, and kidney were dissected, fixed in 4% PFA, and decalcified in 0.5M EDTA/PBS/0.1%PFA. For cryosections, decalcified samples were embedded in OCT/15% sucrose and preserved at −80 °C. Sections (6–12 μm) were cut using a Thermo cryostat and CryoJane tape system. Paraffin-embedded samples were cut into 4–6 μm sections, followed by deparaffinization and heat-mediated antigen retrieval treatment. Multiplex immunofluorescence assay was conducted as previously described (Gu et al., 2020; Ye et al., 2021; Zhang et al., 2020). Briefly, sections were incubated with primary antibody for 2 h, followed by detection using HRP-conjugated secondary antibody and TSA-fluorophores (Histova Biotechnology, NECC7100). Then, the primary and secondary antibodies were eliminated by heating the slides in retrieval/elution buffer (Histova Biotechnology, ABCFR5L) for 10 s at 95 °C using microwaves or by incubating slides in elution buffer (Histova Biotechnology, ABCCC30) for 20–30 min at 37 °C. Serially, each antigen was labeled with distinct antibodies and TSA fluorophores. Multiplex antibody panels applied in this study include: GFP (CST, 2956, 1:400), RFP (Rockland, 600-401-379, 1:800), Pdgfrb (CST, 3169, 1:400), Emcn (ThermoFisher, 14-5851-82, 1:800), Sstr2 (Abcam, ab134152, 1:800), N-cadherin (Abcam, ab76011, 1:500), Pdpn (Sino biological, 50256-R066, 1:1000), CD200 (Novusbio, BAF3355, 1:400), CD51 (Abcam, ab208012, 1:400), Ly51 (Sino biological, 50082-T24, 1:1000), Thy1 (Sino biological, 50461-T44, 1:1000), CD105 (Abcam, ab221675, 1:400), CD31 (CST, 77699, 1:400), CD45 (CST, 70257, 1:400), Ter119 (Biolegend, 116201, 1:1000), Ki67 (CST, 12202, 1:400), Lepr (Novusbio, BAF497, 1:200), Hgs (Abcam, ab155539, 1:300), Txnip (Abcam, ab188865, 1:500), F13a1 (Abcam, ab76105, 1:500), Pla2g10 (Abcam, ab166634, 1:800), and Sox9 (Abcam, ab185966, 1:400). After sequentially detecting all the antibodies, the slides were imaged using a confocal laser scanning microscopy platform Zeiss LSM880 equipped with 405 nm/458 nm/488 nm/514 nm/543 nm/594 nm/633 nm lasers. Some data were further processed and statistically analyzed using the Bitplane Imaris software (Bitplane AG) or the HALO image analysis platform (Indica Labs).

### Histological Analysis

Decalcified or undecalcified samples were embedded in paraffin, cut into 5 μm sections, and subjected to Safranin O and Fast Green staining or von Kossa staining according to the standard protocol.

### Statistical Analysis

Data are expressed as mean ± standard deviation (SD) and significance was set at p < 0.05 (see each figure for details). The sample sizes are shown in the corresponding figure legends. Unpaired one-tailed Student’s t-test was used for two-group comparisons. One-way ANOVA followed by Tukey’s post hoc test was performed for multiple comparisons. All statistical analyses were performed using GraphPad Prism 7 (GraphPad Software).

### Lead contact

Further information and requests for resources and reagents should be directed to and will be fulfilled by the lead contact, Xiao Yang.

### Materials availability

All unique/stable reagents generated in this study are available from the lead contact with a completed Materials Transfer Agreement.

### Data and code availability

The sequencing dataset generated in this study is available in the NCBI Gene Expression Omnibus database (http://www.ncbi.nlm.nih.gov/geo/). Accession number: GSE205945.

## SUPPLEMENTAL INFORMATION TITLES AND LEGENDS

**Figure S1.**
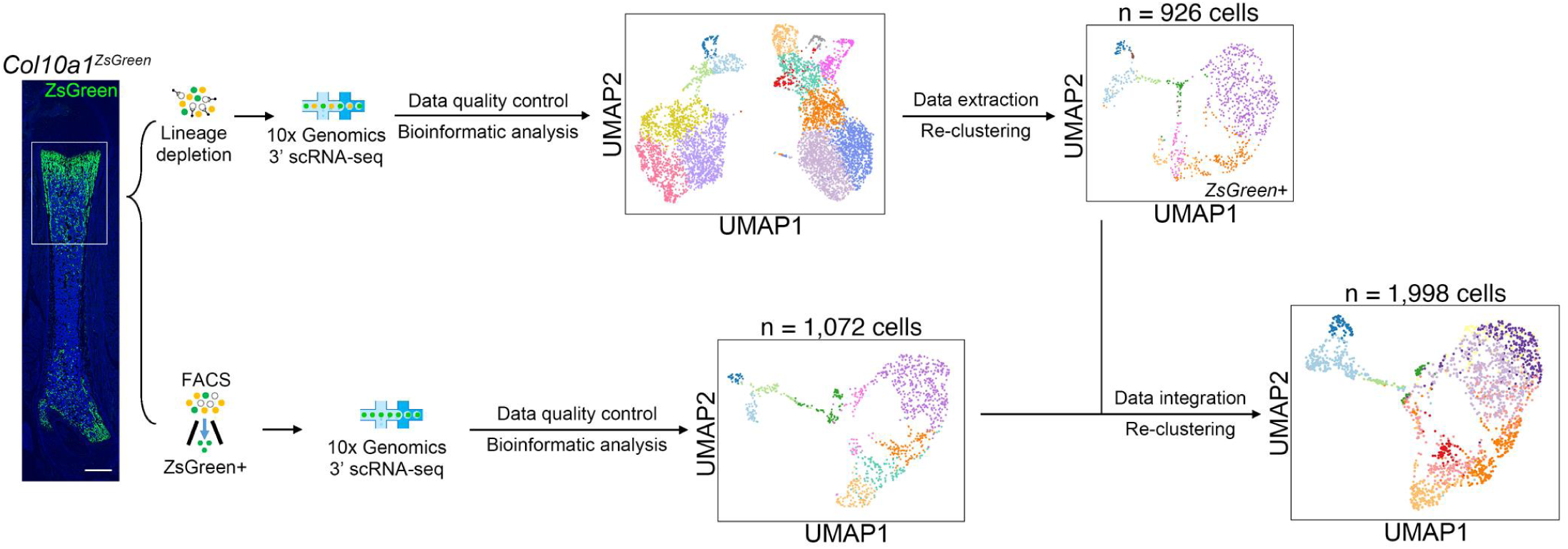
Workflow of Single-cell Analysis of Skeletal Cells, HCs and Descendants of the Developing Bone. Related to Figure 1 and Figure 2. Schematic representation of the strategies to obtain the scRNA-seq data of HCs and their descendants (related to Figure 1C and Figure 2A). The framed part of hindlimb bones from P7.5 *Col10a1^ZsGreen^*mice are used to establish the scRNA-seq transcriptomes. The upper panel shows the technological flow that algorithmically extracts *ZsGreen*-expressing cells from the unsorted scRNA-seq database of skeletal cells. The lower panel shows another method that directly isolates ZsGreen^+^ cells by FACS to establish the scRNA-seq transcriptomics. These scRNA-seq databases are integrated into a new database that is constitutive of HCs and their descendants. They are visualized by uniform manifold approximation and projection (UMAP). Scale bar, 500 μm.

**Figure S2.**
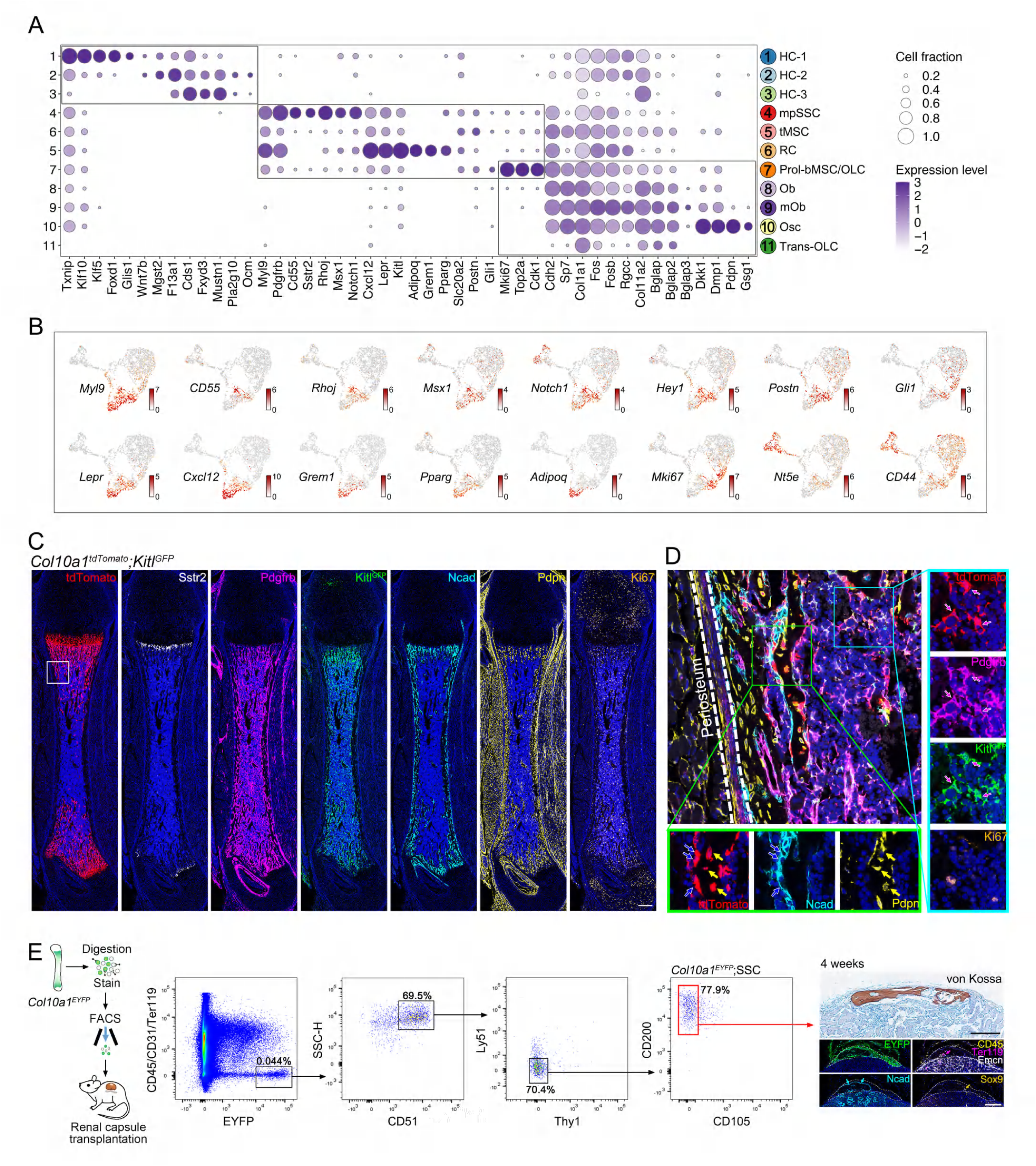
ScRNA-seq and *in situ* Mapping of HCs and Descendants of the Developing Bone. Related to Figure 2. (A) Dot plot shows the expression level of selected cluster-enriched genes in 11 clusters. Dot size indicates the cell fraction of each cluster that expresses listed genes. Purple color intensity indicates scaled average expression level. Three grey-coded frames indicate genes enriched in HC clusters, bMSC clusters, and OLC clusters, respectively. (B) Selected cluster-enriched genes are superimposed on the UMAP plot. Crimson color intensity indicates the Log2 expression level of genes. (C) Representative multicolor IF staining of femur section from P7 *Col10a1-Cre;ROSA26^LSL-tdTomato^;Kitl^GFP^* (*Col10a1^tdTomato^;Kitl^GFP^*) mice (related to Figure 2C). Scale bar, 250 μm. (D) Higher magnification view of the white framed areas that are labeled in (C). Framed areas on the femoral endosteum (green frame) and bone marrow (cyan frame) show where HC-derived Ncad^+^/Pdpn^+^ osteoblasts and Pdgfrb^+^Kitl^high^ reticular cells reside, respectively. (E) HC-derived SSCs form bony organoids. Representative FACS plots of HC-derived SSCs (*Col10a1^EYFP^*;SSC) that are used for renal capsule transplantation. The representative von Kossa and multicolor IF staining indicate that bony organoids formed by the HC-derived SSCs consist of Sox9^+^ chondrocytes, Ncad+ osteoblasts, as well as host-derived blood vessels and hematopoietic cells. Dashed lines indicate the margins of the EYFP^+^ bony organoids. Scale bar, 200 μm.

**Figure S3.**
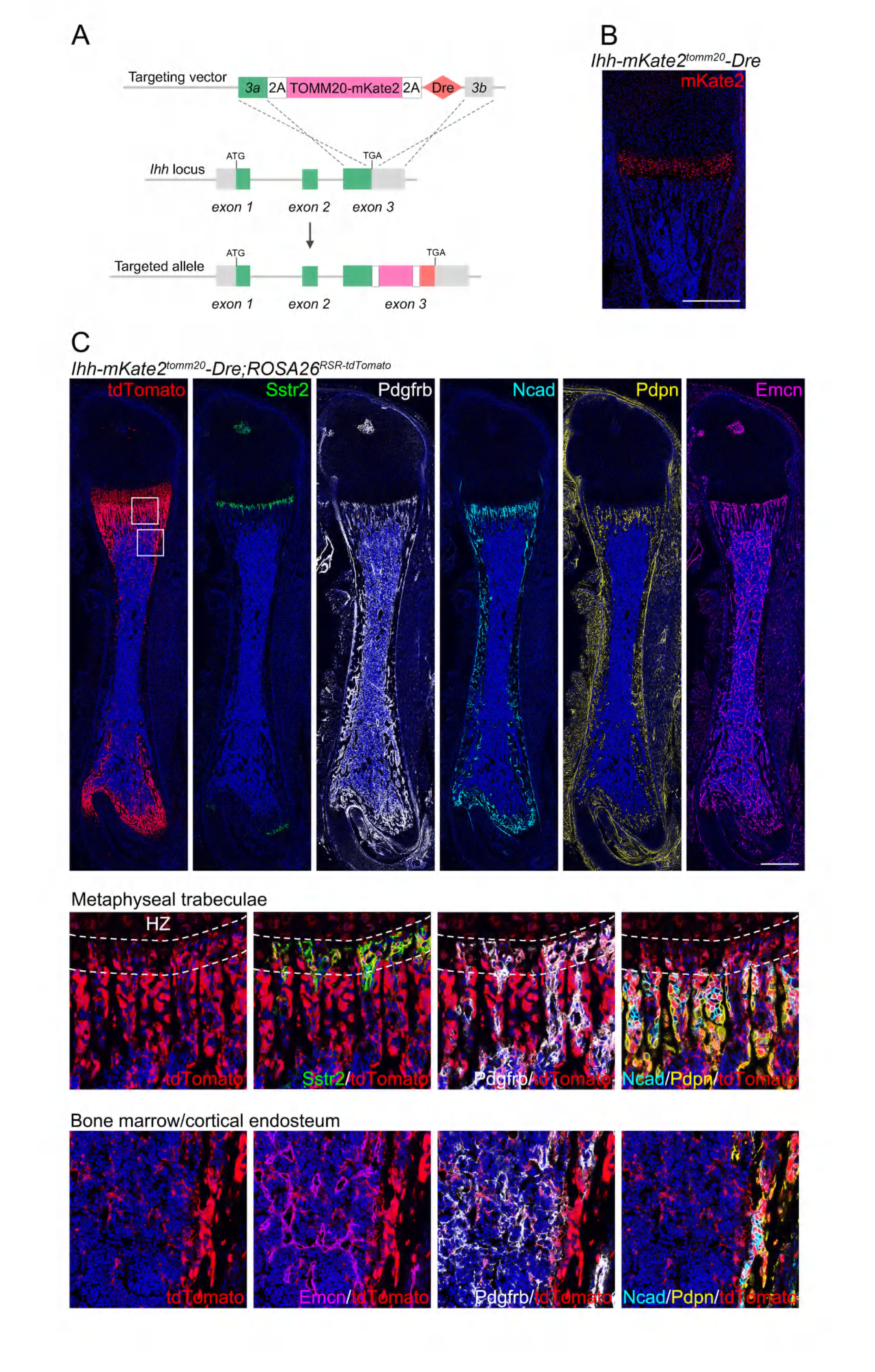
*In situ* Mapping of Pre-HCs and Descendants in *Ihh-mKate2^tomm20^- Dre;ROSA26^RSR-tdTomato^* Mice. (A) Strategy generating *Ihh-mKate2^tomm20^-Dre* knock-in alleles by CRISPR-Cas9 technology. (B) Representative IF staining of mKate2 fluorescent protein in femur section from P7 *Ihh-mKate2^tomm20^-Dre* mice. Mitochondrion-targeted mKate2 is presented in pre-HCs and HCs. Scale bar, 500 μm. (C) Representative multicolor IF staining of femur section from P8 *Ihh-mKate2^tomm20^- Dre;ROSA26^RSR-tdTomato^* mice. Higher magnification views of the white framed areas are shown below, which demonstrate pre-HC/HC-derived Sstr2^+^ mpSSCs, Pdgfrb^+^ RCs, Ncad^+^/Pdpn^+^ OLCs at the metaphyseal trabeculae, bone marrow and cortical endosteum, respectively. The upper metaphyseal region right beneath the hypertrophic zone (HZ) is framed by dashed lines. Scale bar, 500 μm.

**Figure S4.**
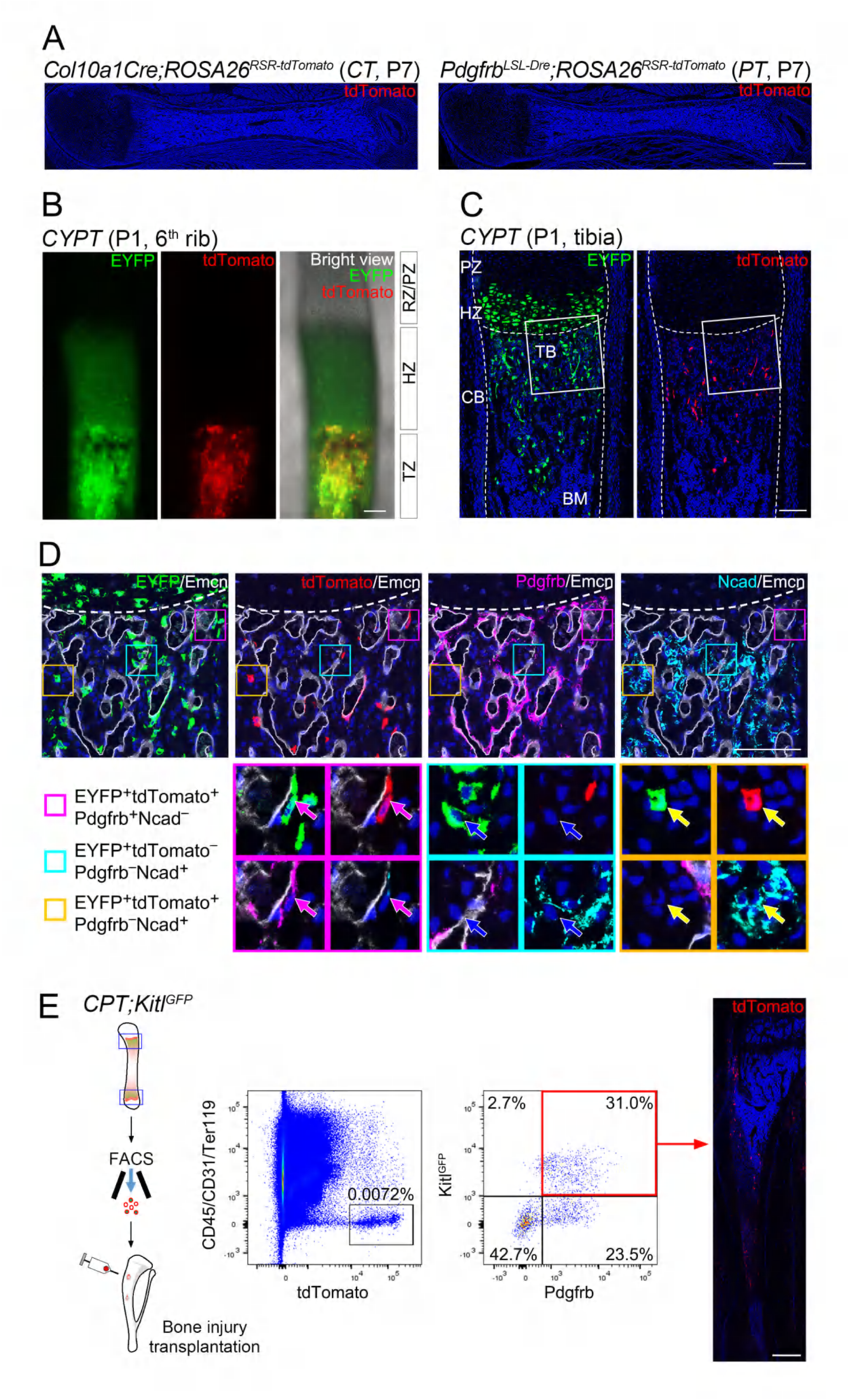
Supplementary Data Related to Figure 3. (A) tdTomato expression in P7 *Col10a1Cre;ROSA26^RSR-tdTomato^*(*CT*) and *Pdgfrb^LSL-Dre^;ROSA26^RSR-tdTomato^*(*PT*) mice verified no cross-reaction of Cre on Rox as well as efficient transcriptional blocking of STOP sequence in *Pdgfrb^LSL-Dre^*line, respectively. Scale bars, 500 μm. (B) Whole-mount fluorescent imaging shows tdTomato and EYFP expression in the 6^th^ rib of the P1 *CYPT* mouse. Scale bar, 100 μm. (C, D) Representative multicolor IF staining of P1 *CYPT* tibia. Dashed lines indicate the outline of the tibia and the chondro-osseous junction of the growth plate. RZ, resting zone; PZ, proliferating zone; HZ, hypertrophic zone; TB, trabecular bone; CB, cortical bone; BM, bone marrow. Scale bars, 100 μm. White framed areas of the chondro-osseous junction are shown as (D) at higher magnifications. The magenta, cyan, and yellow framed areas and arrows show the EYFP^+^tdTomato^+^Pdgfrb^+^Ncad^−^ mesenchymal cells, EYFP^+^tdTomato^−^Pdgfrb^−^Ncad^+^ osteoblasts, and EYFP^+^tdTomato^+^Pdgfrb^−^Ncad^+^ osteoblasts, respectively. Scale bars, 100 μm. (E) Representative experimental results for sorting and characterization of primary tdTomato^+^Pdgfrb^+^Kitl^high^ RCs (related to Figure 3H and Figure 3I). The representative IF staining shows the recipient tibia 14 days after tdTomato^+^Pdgfrb^+^Kitl^high^ RCs transplantation. Scale bars, 500 μm.

**Figure S5.**
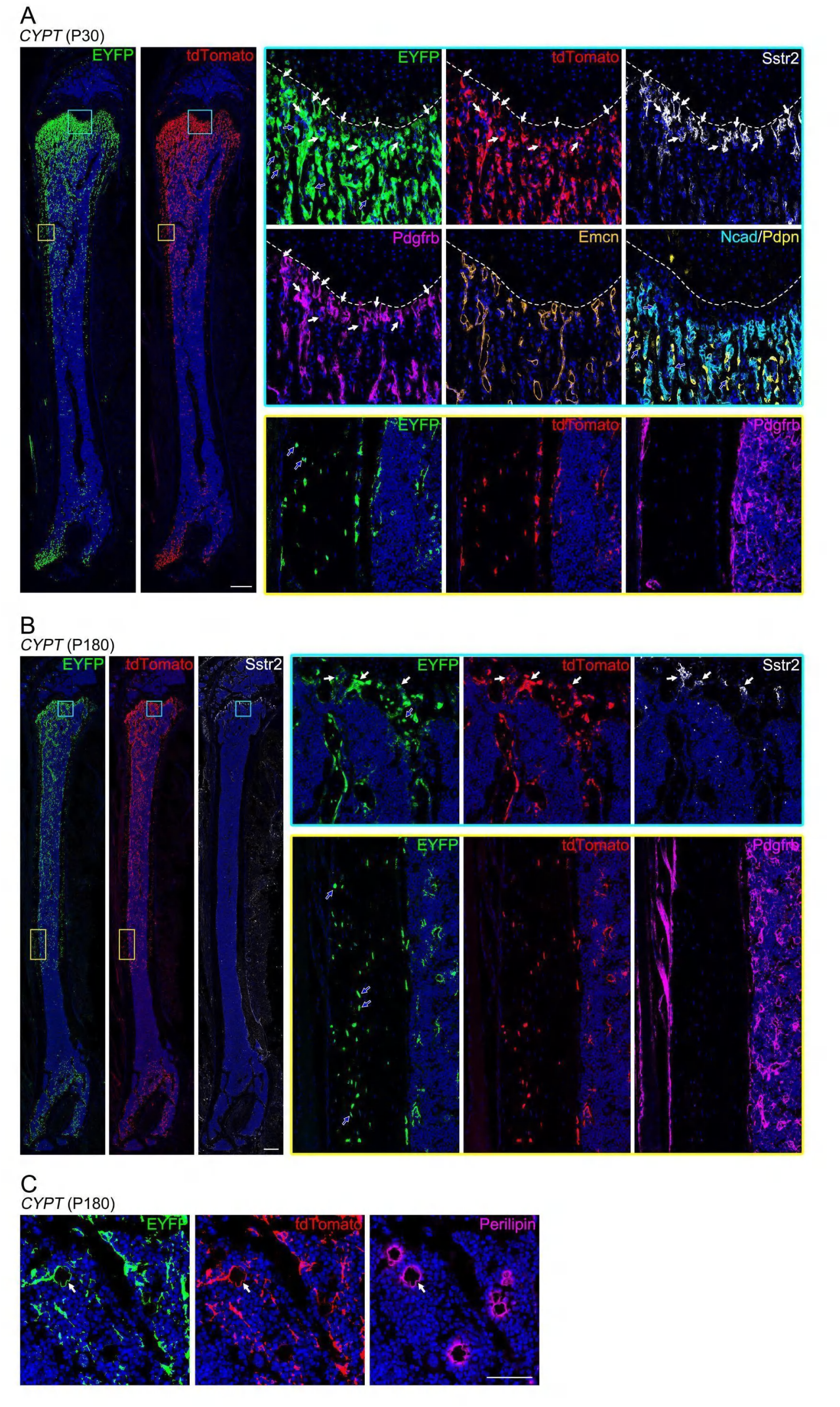
Mapping Two-fated HC Descendants in Adult Mice. Related to Figure 4. (A, B) Representative multicolor IF staining maps HC-derived, two-fated cell compartments in femur sections from P30 and P180 *CYPT* mice. Cyan- and yellow-framed areas indicate the metaphyseal trabeculae, the diaphyseal cortical bone, and bone marrow, and are shown adjacently at higher magnifications, respectively. Dashed lines indicate the chondro-osseous junction of the growth plate. white arrows: EYFP^+^tdTomato^+^Pdgfrb^high^Sstr2^+^ mpSSCs; blue arrows: EYFP^+^tdTomato^−^ OLCs embedded in bone. Scale bar, 500 μm. (C) Representative multicolor IF staining shows adipogenesis within the EYFP^+^tdTomato^+^ compartment in the bone marrow of P180 *CYPT* mice. Scale bar, 50 μm.

## Notes

### Competing Interest Statement

The authors have declared no competing interest.

